# Hippocampal representations of alternative possibilities are flexibly generated to meet cognitive demands

**DOI:** 10.1101/2024.09.23.613567

**Authors:** Alison E. Comrie, Emily J. Monroe, Ari E. Kahn, Eric L. Denovellis, Abhilasha Joshi, Jennifer A. Guidera, Timothy A. Krausz, Joshua D. Berke, Nathaniel D. Daw, Loren M. Frank

**Affiliations:** Neuroscience Graduate Program, University of California San Francisco; San Francisco, CA 94158, USA; Department of Physiology and Psychiatry, University of California, San Francisco; San Francisco, CA 94158, USA; Princeton Neuroscience Institute, Princeton University; Princeton, NJ 08544, USA; Howard Hughes Medical Institute; Chevy Chase, MD 20815, USA; Medical Scientist Training Program, University of California, San Francisco, San Francisco, CA 94143, USA; Kavli Institute for Fundamental Neuroscience, University of California, San Francisco; San Francisco, CA 94158, USA; Department of Neurology and Department of Psychiatry and Behavioral Science, and Weill Institute for Neurosciences, University of California, San Francisco, San Francisco, CA 94158, USA; Department of Psychology, Princeton University; Princeton, NJ 08544, USA

**Keywords:** hippocampus, learning, memory, spatial navigation, value-based decision making, updating, internal model, foraging, theta sequences, cognitive map

## Abstract

The cognitive ability to go beyond the present to consider alternative possibilities, including potential futures and counterfactual pasts, can support adaptive decision making. Complex and changing real-world environments, however, have many possible alternatives. Whether and how the brain can select among them to represent alternatives that meet current cognitive needs remains unknown. We therefore examined neural representations of alternative spatial locations in the rat hippocampus during navigation in a complex patch foraging environment with changing reward probabilities. We found representations of multiple alternatives along paths ahead and behind the animal, including in distant alternative patches. Critically, these representations were modulated in distinct patterns across successive trials: alternative paths were represented proportionate to their evolving relative value and predicted subsequent decisions, whereas distant alternatives were prevalent during value updating. These results demonstrate that the brain modulates the generation of alternative possibilities in patterns that meet changing cognitive needs for adaptive behavior.

## Introduction

Animals are continually faced with decisions about what to do and where to go next. In the context of behavioral tasks, a long tradition of animal experiments and behavioral models suggest that the brain can make adaptive decisions by comparing the expected values of available options, which are learned through experience. As an example, in spatial settings, an animal may compare the expected values of different rewarded locations and associated paths. After making a choice, a rewarding outcome would lead to an update that increases the stored value of the rewarded location and the path taken to get there, enabling subsequent adaptive decisions even as outcomes change^1–4^.

While the idea of retrieving and updating expected values of possible options seems relatively simple, in real-world situations like navigation, choices often lead to outcomes that are distant in space or time. This poses a challenge: the brain must go beyond current experience to decide among or learn about alternative “non-local” possibilities. For instance, when making a choice among nearby routes, the brain may retrieve values related to their ultimate destinations. Further, in complex structured scenarios, rewards received in one location can imply information about the availability of rewards in other places, as in zero-sum situations or multi-step planning tasks. In such cases, the brain may take advantage of learned structure to make inferences across space, and update not only the value of the current location but also the values of alternative locations and paths^5–8^. Additionally, unlike many laboratory tasks, naturalistic scenarios can have many alternatives available at once^9–14^. This indicates a need to prioritize^15^; when deciding among or updating different possibilities, some judicious mechanism is required to consider the most relevant non-local alternatives to meet current demands.

The neural mechanisms that enable such prioritized computations about relevant non-local possibilities during complex behavior are not understood. Existing data indicate the hippocampus and its representations of space as a starting point. First, the hippocampus is critical for rapid learning and performance of spatial tasks where animals must learn the locations of and routes among rewarded locations^16–23^. Second, the hippocampus is well known for spatially tuned “place cells” whose activity typically signals the actual location of the animal^24^. These cells are often described as the substrate for a cognitive map, or an internal model of the world, that encodes the relationships among both locations and experiences more broadly^25–27^. Critically, while place cells are best known for coding an animal’s actual position, they are also capable of expressing non-local representations of alternative locations at a sub-second timescale^8,28–34^. Finally, natural behavior often involves experience-guided decision making during active navigation, and hippocampal non-local representations can be regularly expressed during movement^35,36^. Thus, during active behavior, generating a non-local representation corresponding to a particular alternative location could serve to retrieve or update information associated with that location, including its value.

Non-local representations have been linked to cognitive processes for both decision making and learning during navigation. These population-level representations are most often associated with the sequential firing of place cells corresponding to a trajectory through locations behind, at, and ahead of the animal’s actual location^28,37,29,38,39^. In the context of decision making, as animals approach a choice point, these representations can sweep along future paths ahead, which is evocative of a role in retrieving at least immediately upcoming options^10,35,40–46^. These sequences also engage place cell activity on timescales consistent with synaptic plasticity, suggesting a role in learning^28,29,47–51^, and potentially updating internal representations based on experience.

Yet, whether or how the brain generates representations of different alternatives as cognitive demands change throughout experience-guided decision making and learning remains unclear. We therefore combined approaches typically used to separately study decision making and reinforcement learning, or experience-guided navigation. We developed a dynamic patch-foraging task where changing reward probabilities across six locations challenged animals to continually update their internal models to make experience-guided choices about where to go for reward. By leveraging a computational model, we estimated internal cognitive variables from animal behavior related to both value-guided decision making and value updating. As these cognitive variables evolved across successive trials of experience, we monitored hippocampal neural activity to identify non-local representations expressed during active navigation. We observed a range of representations of alternatives, corresponding to potential paths not only ahead of, but also behind the animal, including along counterfactuals, and to distant locations in remote foraging patches. We further found that these representations were modulated across successive trials in two distinct patterns, one for representational content and the other for spatial extent, each of which was related to distinctly evolving cognitive variables. These findings demonstrate that mechanisms exist that regulate the expression of distinct non-local possibilities in conjunction with cognitive needs.

## Results

### Rats make experience-guided decisions in the Spatial Bandit task

We developed a dynamic foraging task where performance could benefit from representing alternative possibilities, both for deliberating among alternatives and for updating information about alternatives. This “spatial bandit” task combined features of spatial memory and decision-making paradigms (Fig. 1A). First, as in classic spatial memory behaviors, rats (n=5) navigated a maze based on prior experience to reach reward locations. The track was made up of three Y shaped “foraging patches” radiating from the center of the maze, and each patch contained two reward ports at the ends of the linear segments (Fig. 1A, left). The multiple bifurcations and reward locations provided many opportunities for deliberation among alternative options and reward-based updating. Second, as in classic decision-making tasks, we introduced uncertainty by dispensing rewards probabilistically. Each port was assigned a nominal probability of reward, p(R), of 0.2, 0.5, or 0.8 (Fig. 1A), and one of the three patches had a greater average p(R) than the other two. On any visit to a port the rats either did or did not receive reward, determined stochastically by the p(R), and consecutive visits to the same port were never rewarded. Thus, animals could only infer each port’s reward probability state through experiences across multiple visits, and because rewards were delivered probabilistically, it could take many trials for an animal’s experience-based estimated p(R) to converge to the true value.

**Figure 1.**
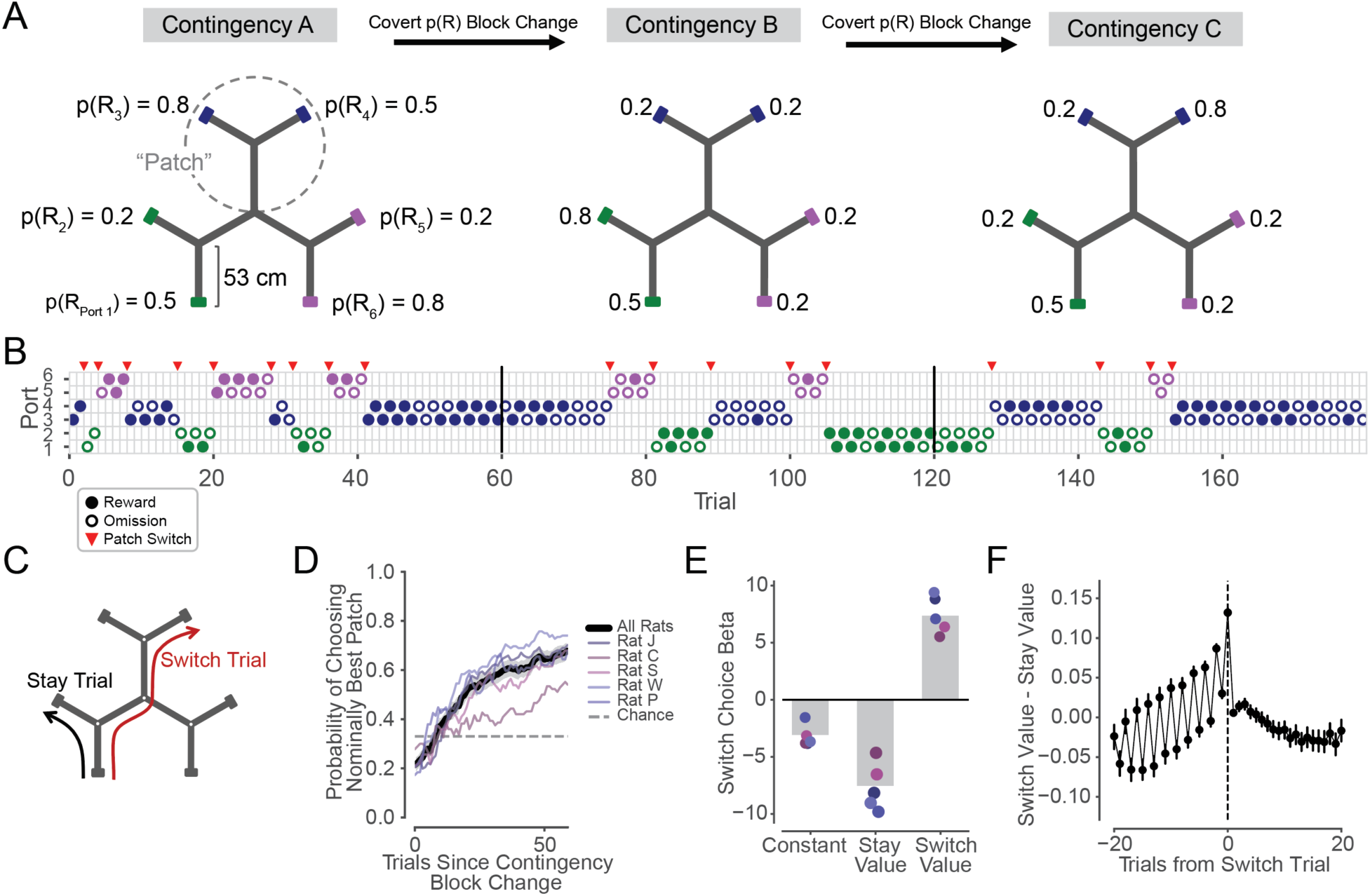
Rats make experience-guided decisions in the Spatial Bandit task. **(A)** Track schematic with example reward Contingencies A-C from one behavioral epoch. The track contains three Y-shaped “patches,” each with 2 reward ports located at the ends of linear segments. Probability of reward at each of 6 identical reward ports is indicated as p(R_1-6_). Ports are colored by patch for figure visualization only. Uncued reward contingency changes occur every 60 trials. **(B)** Example sequence of choices from one behavioral session. Vertical black lines indicate contingency changes shown in **A**. Circles indicate chosen port on each trial and are colored by patch. Filled circles represent rewarded trials, empty circles represent unrewarded trials. Red triangles indicate a patch Switch. **(C)** Schematic of example Stay (black) and Switch (red) trial trajectories. White dots at track segment intersections indicate choice points. **(D)** Proportion of trials in which the animal chose a port in the patch with the highest average nominal reward probabilities, as a function of trial number within the contingency block. Black line and grey shading indicate average across rats and 95% CI on the mean. Distribution on 60th trial of blocks is significantly different from chance level of 1/3 in each animal (n=168, 87, 78, 131, 87 blocks for animals J, C, S, W, and P, respectively, p=2.7e-22, 8.7e-5, 6.4e-10, 2.5e-21, 6.1e-11, binomial test), and distribution on 60th trial is also significantly different from values on first trials of blocks (p=9.6e-20, 3.9e-4, 2.5e-9, 3.2e-18, 1.5e-11, Z test for proportions) **(E)** Regression coefficients on model-estimated values of Stay locations and Switch locations for the prediction of Stay or Switch choice on each trial. All individual coefficients on the Stay and Switch values are significantly different than zero, indicating that rats rely on reward history from both current “Stay” patch and alternative “Switch” patches to make Stay or Switch choices (for each animal, p_Constant_=2.1e-5, 2.3e-16,4.1e-3, 5.0e-3, 3.3e-1, p_Stay_=4.9e-85, 3.2e-40, 2.4e-42, 2.7e-76, 4.4e-70, p_Switch_=7.2e-7, 2.0e-10, 3.0e-3, 2.7e-4, 2.7e-2, from n = 5416, 8572, 9608, 7450, 4732 Stay trials and n = 344, 398, 583, 467, 488 Switch trials). Grey bars indicate averages and coefficient points are colored by subject per legend in **D**. **(F)** Relative value of Switching versus Staying increases across trials leading up to patch Switch choices, peaks on Switch trials, and decreases following patch Switch trials. Switch trial values are statistically greater than Stay trial values in the 20 trials before (p=1.1e-52, 1.3e-42, 6.9e-117, 5.2e-115, 1.9e-111, Wilcoxon rank sum test) and after (p=9.2e-69, 1.3e-57, 1.5e-143, 2.7e-139, 3.2e-131) Switch trials in each individual animal. Data plotted are from all animals; error bars show 95% CI on the mean, and individual animal data is shown in Fig. S5E.

During run sessions, reward probabilities around the track covertly changed in blocks (as in Fig. 1A,B) every 60 (n=4 animals) or 80 (n=1 animal) trials. Each day of task experience consisted of up to 8 run sessions, separated by rest sessions in a rest box, and each run session consisted of 180 (n=4 animals) or 160 (n=1 animal) trials. A trial was defined as the period between a departure from one port to a departure at another port, and animals individually completed 5760, 8970, 10191, 7917, and 5220 trials, providing a large dataset compatible with behavioral and neural analyses. Importantly, the reward contingency block transitions were uncued and changed which patch was associated with the highest overall nominal probability of reward. These reward probability transitions thereby encouraged experience-based, adaptive decision making.

Animals exhibited choice behavior akin to patch foraging^52^ (Fig. 1B). They began by serially exploring the ports in the three patches, often making repeated choices to alternate between ports within a patch (Stay trials) and occasionally making flexible choices to navigate to an alternative patch (Switch trials) (Fig. 1B,C, Fig. S1A). On average, animals were more likely to make Switch choices in the first 20 versus final 20 trials of each contingency block (each rat p<0.01, Wilcoxon rank sum test). Animals adapted their choices based on their dynamic reward experiences across reward ports, trials, and contingencies (as in Fig. 1B). As a result, animals generally learned and remained within the patch with the highest nominal reward probabilities by the end of each block (Fig. 1D).

The patterns of these Stay and Switch choices indicated a decision strategy that used reward history information related to both the current patch and to distant alternative patches. To understand how reward information related to animals’ choices at a single-trial level, we fit a statistical behavioral learning model to each animal’s sequence of port choices and reward outcomes (see Methods). Briefly, the model estimates a belief about the value of each port on each trial based on the reward outcomes experienced at each port. The model implements a Bayesian version of recency-weighted reward learning^15^, which is analogous to Q-learning, but estimates reward values with a Beta distribution that updates based on the binary outcomes observed at each Bernoulli reward port (Fig. S5A-C). In addition, the model accounts for the possibility of forgetting when not visiting a port by allowing a slow flattening of an unvisited port’s estimated reward probability distribution toward uniform, where all probabilities of reward were equally likely. This approach enabled us to quantitatively infer from animal behavior the expected values of the animal’s two potential options on each trial: the value of Staying within the current patch or the value of Switching between patches.

Consistent with animals weighing the values of both more local and more remote alternative options, logistic regressions predicting Stay or Switch choices revealed significant effects of not only the Stay value but also the Switch value in each animal (Fig. 1E). The coefficients were negative for Stay values, indicating that animals were less likely to make Switch choices as the value of Staying increased (Fig. 1E). In contrast, the coefficients were positive for Switch values, indicating that animals were more likely to choose to Switch as the value of Switching increased (Fig. 1E). Thus, animals’ behavior depended on previous reward experiences both from nearby Stay locations and more spatially distant Switch locations across the maze.

Since Switch choices were value-guided and often punctuated longer, stable bouts of Stay choices (Fig. 1B,C, Fig. S1A), we anchored further analyses to these self-paced patch changes. As our analyses identified contributions of both local and remote value, we focused on the corresponding decision variable: the relative value of these two options on each trial (Switch value - Stay value; Fig. 1F). To understand how this relative value of Switching versus Staying evolved across trials around animal’s choices to Switch between patches, we computed this decision variable from the behavioral model that estimated these values from animals’ experienced reward histories (rather than using the hidden nominal reward probabilities assigned to each port).

We found that the relative value of Switching versus Staying was low during trials far from a Switch and on average increased over the course of the Stay trials leading up to a Switch choice. On the Switch trial, relative value was highest and then decreased again across trials after the Switch (Fig. 1F). Note that these relative values do not incorporate the bias to Stay (or cost to Switch) as illustrated by the negative constants in Fig. 1E. Importantly, the overall across-trial pattern of increasing and decreasing relative value around Switches reflects a gradual updating of values over successive trial outcomes. This suggests that animals may similarly access internal estimates of values associated with Switch and Stay options when deciding whether to leave the current patch and, after Switching, whether to remain in the new patch or again navigate elsewhere.

We also noted the relative value fluctuations on alternating trials before a Switch (Fig. 1F), which are consistent with animals tending to Switch after visiting the higher value port within a patch. Specifically, when the two ports within a patch had distinct nominal values, animals initiated 68.3% of Switch trials from the higher value port. Thus, the average pattern of relative values across even and odd trials preceding a Switch is consistent with a tendency to alternate between the worse and better ports before departing a patch.

Taken together, these findings provide evidence that animals learn from their changing reward experiences across the maze and leverage this experience to make value-guided decisions among alternative paths. Furthermore, they suggest that seemingly isolated and behaviorally overt Switch choices occur in the context of gradually changing covert reward expectancies that place evolving cognitive demands on the animal across successive trials.

### Representations of alternative paths ahead of the animal

Given these behavioral results, we then asked whether the content of hippocampal representations of alternative paths was also modulated around Switch choices. As previous work has identified non-local representations consistent with possible future locations^35,40–43^, we began by examining non-local representations extending ahead of the animal’s actual position that occurred while the animal was in the first segment of each trial. In this period, the animal approached the first choice point, which is associated with the choice to Stay or Switch. We used an established state space decoding algorithm^53,54^ (2 cm spatial bins, 2 ms temporal bins, see Methods) to assess the instantaneous hippocampal representation of space^55–57^ during navigation (animal speed > 10 cm/s) across all five rats, each implanted with tetrode microdrives targeting the CA1 region of the hippocampus (Fig. S1). We limited our analyses of non-local representations to periods with high confidence that the decoded hippocampal representation was in a non-local track segment, defined as a segment distinct from the one corresponding to the rat’s actual location (Fig. 2A, see Methods).

**Figure 2.**
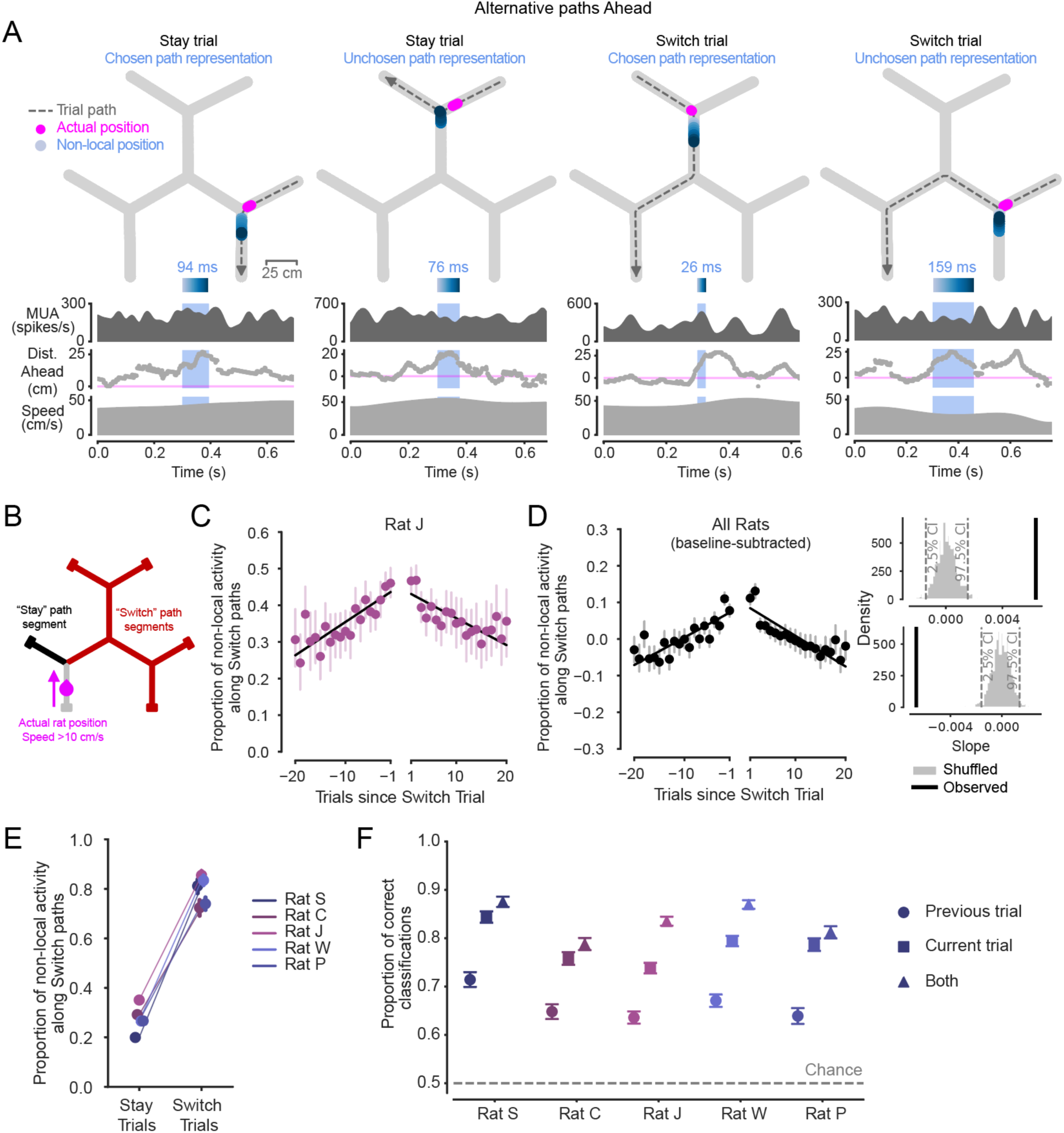
Non-local representations of alternative paths Ahead are enriched across trials before and after patch Switching. **(A)** Examples of non-local representations of alternative paths ahead on Stay and Switch trials. As the animal approaches the choice point, the hippocampus can represent the Stay or Switch path ahead. Top: Animal’s actual position (magenta) and decoded position from hippocampal spiking (blue, shaded by time) during the light blue-highlighted period (below) are plotted on the track. Dashed grey line with arrow indicates animal’s path and direction on the current trial. Highlighted non-local duration is labeled above blue time-shaded heatmap. Top-middle: multiunit spike rate from hippocampal tetrodes before, during, and after highlighted non-local representation. Bottom-middle: Distance from actual position to decoded position, with positive values corresponding to in front of the animal’s heading, and negative numbers corresponding to behind the animal’s heading. Note that blue non-local periods are required to have representations in a different track segment from the animal’s actual position. Bottom: Animal speed. **(B)** Schematic defining Stay and Switch non-local activity ahead, analyzed in **C-F**. Non-local activity was analyzed from times in which the animal was running >10 cm/s from the reward port to the first choice point in the trial (magenta) by assessing the per-trial proportion of time in which non-local activity extended along the path ahead consistent with Switching (red) out of all non-local activity times along both the Stay (black) and Switch (red) paths. Here, the relevant choice point is ahead of the animal at the intersection between the grey, black, and red track segments. **(C)** Proportion of all non-local activity that represents paths consistent with Switching during the approach of the first choice point across Stay trials preceding and following Switch trials in one rat. Left and right x axes separated to reflect patch change. Error bars are 95% CIs on the mean. Linear regressions show increasing and decreasing proportions across trials before and after patch Switch trials. **(D)** Left: Proportion of all non-local activity that represents paths consistent with Switching during the approach of the first choice point across all animals (n=5 rats), normalized per animal by subtracting off the average baseline proportion on Stay trials. Error bars are 95% CIs on the mean. Pre- and post-Switch linear regressions overlaid. Upper right: observed slope of pre-Switch regression (black) is significantly different from 0 (p<0.002) based on 1000 shuffles of the underlying data (grey). Lower right: observed slope of post-Switch regression (black) is significantly different from 0 (p<0.002) based on 1000 shuffles of the underlying data (grey). Both slopes were individually significant in all five animals (Fig. S2A). **(E)** Proportion of all non-local activity that represents paths consistent with Switching during the approach of the first choice point in each animal on Stay trials and Switch trials. Error bars are 95% CIs on the mean. The proportion on Switch trials is greater than on Stay trials for all animals (p=2.7e-192, 4.2e-108, 1.0e-100, 2.1e-315, 1.6e-201, Wilcoxon rank sum test, for n=313, 335, 523, 422, 452 Switch trials, and n=3349, 3803, 5600, 5028, 3294 Stay trials). **(F)** Cross-validated accuracy of logistic regressions that predicted Stay or Switch choices based on the proportion of all non-local activity that represents paths consistent with Switching during the approach of the first choice point. Neural data from each of three trial types: previous trial (circle marker), current trial (square marker), or both (triangle marker). Error bars are 95% CIs on the proportion. Training data were balanced, so chance level was 0.5. Accuracy of model using previous trial neural data is significantly greater than chance level in all animals (p=3.2e-63, 1.1e-43, 4.0e-133, 9.7e-131, 3.3e-60, Z test for proportions), accuracy from model using current trial neural data is greater than accuracy from model using previous trial neural data (p=2.3e-39, 7.0e-27, 6.0e-33, 2.8e-46, 6.3e-43), and accuracy of model based on both trials is greater than accuracy of current trial model (p=1.1e-4, 1.2e-3, 4.3e-39, 4.9e-26, 5.6e-3).

We observed non-local representations that reflected locations along either of the two paths ahead of the choice point (Fig. 2A, Supplemental Video 1). We also verified the expected organization of these non-local representations of paths ahead based on the phase of the hippocampal theta rhythm^28,29,37,58,59^ (Fig. S2C). On each trial, we then classified non-local representations as reflecting paths that, if physically traversed, would either lead the animal to Stay in the patch or to Switch between patches (Fig. 2B). We began by examining the Stay trials leading up to and following Switch choices. Critically, this enabled us to assess the content of non-local representations as the relative value of Stay and Switch paths changed (Fig. 1F), while the behavioral choice to Stay remained constant.

Strikingly, the relative representation of Switch and Stay paths mirrored their evolving relative value. Across trials preceding a patch Switch, non-local representations of Switch paths ahead became increasingly prevalent relative to Stay paths (Fig. 2C,D). All animals exhibited this pattern, showing an approximately 1.3-1.5 fold increase in the likelihood of non-local representations consistent with the unchosen Switch versus the chosen Stay paths over the 20 trials before a Switch choice (Fig. 2C,D, Fig. S2A). This culminated in a ratio approaching equal (0.5) representation of each path ahead on the trial before a Switch trial (Fig. 2C, Fig. S2A). Additionally, on Stay trials after the animal arrived in a different patch following a Switch, non-local representations were again enriched for the unchosen Switch path. Then, the longer the animal chose to Stay, the less the non-local representations reflected the Switch path (Fig. 2C,D, Fig. S2A).

These increases and decreases in the proportion of Switch path representations were driven largely by changes in the amount of time spent representing the Switch path, while the level of Stay path representation remained, on average, relatively stable across successive Stay trials (Fig. S3A). We also confirmed that the approximately symmetrical increasing and decreasing pattern around a Switch was not driven by periods when the animal and decoded non-local spatial representation were very close by, as is possible near choice points, by requiring representations to extend at least 10 cm from the animal (Fig. S3C). Additionally, this pattern was not driven only by short Stay bouts but was also seen when analyses were restricted to long bouts (Fig. S3F). Further, these patterns could not be attributed solely to differences in head orientation. As expected from prior work^35,40^, we found a weak correlation between head orientation and subsequent Stay or Switch choice, but this was present during both non-local (each rat p<0.01, circular-linear correlation test) and local representations (each rat p<0.01), and we observed that the head direction opposed the non-local content direction on 38.6% of non-local times on average. Similarly, no consistent relationships between running speed and non-local content were observed.

We also ruled out the possibility that low spike rates drove the pattern of representations associated with Switching. No animals showed lower spike rates on Switch than Stay trials (all p>0.05, Wilcoxon rank sum test) and only one animal showed significant non-zero slopes of linear regressions before and after Switches when compared to 1000 shuffles of the underlying data; and, in this case, the slopes increased rather than decreased around Switch trials. Finally, we confirmed that the results were consistent when analyses were limited to continuous-state decoding periods (see Methods).

Notably, the modulation of non-local representations of paths ahead of the animal before and after Switch trials resembles the changes in relative value between the Switch and Stay options (Fig. 1F). That is, trials with a higher relative value of Switching were associated with greater relative representation of Switch paths. This same relationship could also explain differences in non-local representations across Stay and Switch trials. Stay trials were on average associated with approximately 20-30% non-local representation of paths consistent with Switching, whereas the complementary bias was seen on Switch trials, in which 70-80% of the alternative representations were of the Switch path (Fig. 2E).

To quantitatively assess the contributions of underlying values and behavior to the content of non-local representations on Stay trials, we built generalized linear models and asked how the proportion of non-local activity along Switch versus Stay paths varied with the relative value of Switching, and the number of trials away from the next behavioral Switch choice. Both relative Switch value and number of trials until the Switch significantly contributed in each animal (β_value_ = 1.47, 1.01, 0.89, 0.93, 1.51, with p-values = 1.45e-06, 2.29e-06, 5.25e-07, 2.77e-05, 1.35e-06, and β_behavior_ = -0.0065, -0.0067, -0.0049, -0.010, -0.0097, with p-values = 1.65e-02, 1.18e-05, 8.62e-04, 1.25e-06, 3.75e-04). Interestingly, this relationship was not present on the Switch trials themselves, when there was not a consistently significant contribution of relative value to the proportion of non-local representation along Switch paths. This indicates that non-local content evolves across Stay trials as a function of both the changing expected value of Switching versus staying and the behavioral number of trials from Switching, and that upon Switch trials, non-local content primarily reflects the choice to Switch.

Together, these findings (Fig. 2A-E) indicate that the hippocampus continually tunes the retrieval of spatial alternatives in a pattern related to their evolving relative value. In the context of experience-guided decision making, our results are consistent with an across-trial process where internal sampling of alternatives is biased by relative expected value, such that as these estimated values become more similar—and the cognitive demand for distinguishing between them potentially greater—the options are sampled more equally. Recruiting representations of relevant paths based on their viability for making the best choice, in turn, could enable more accurate comparisons of the values^60^.

The possibility that non-local representations are engaged across trials in an internal sampling process led us to ask whether the proportion of non-local representation of the Switch path was predictive of the future choice to Stay or Switch. We reasoned that proportionally more representation of the Switch option on the Stay trial an entire trial in advance of the Switch could provide samples of the relatively high value Switch option (Fig. 1F) that in turn could influence the decision on the next trial. To investigate this at the level of single trials, we used a cross-validated logistic regression to predict whether an animal would choose to Stay or Switch on each trial based on the proportion of non-local representation corresponding to Switch paths as the animal approached the first choice point of each trial. This model predictor, the proportion of non-local Switch path representation, corresponds to the same metric as in Figs. 2C-E. Training data were balanced such that chance level was 0.5.

We found evidence that non-local representations could be used in an across-trial decision process. Non-local representations occurring on the previous trial predicted the subsequent trial’s Stay or Switch choice better than chance. This indicates that the increased alternative representation on the Stay trial before a Switch trial was predictive of a future Switch choice an entire trial in advance, even when the immediately upcoming choice on the current trial was still to Stay (Fig. 2F). This choice prediction was even more accurate when considering only the non-local representations that occurred during the approach of the choice point on the current trial (Fig. 2F), which is expected from Fig. 2E. Strikingly, including non-local representations on both the previous trial and the current trial led to an even better prediction of the choice the animal would make on the current trial (Fig. 2F). While Fig. 2C-E indicate a ramping process on average across bouts of Stay trials before patch Switches, this regression result extends the observation of an across-trial modulation to the level of individual choices within bouts.

### Representations of alternative paths behind the animal

The idea that representations of alternatives may be flexibly generated to meet decision-making needs across trials, rather than only for the immediate future choice, led us to ask next whether non-local representations of alternatives could be expressed at times when there was no immediately upcoming choice point. Here we focused on periods when the animal was traversing the final track segment of a trial and approaching a reward port. During this period, we found that non-local representations not only corresponded to the path the animal recently traversed, but also the unchosen Stay or Switch path that the animal could have come from or gone to but did not. This is consistent with the representation of counterfactuals (Fig. 3A,B, Supplemental Video 2).

**Figure 3.**
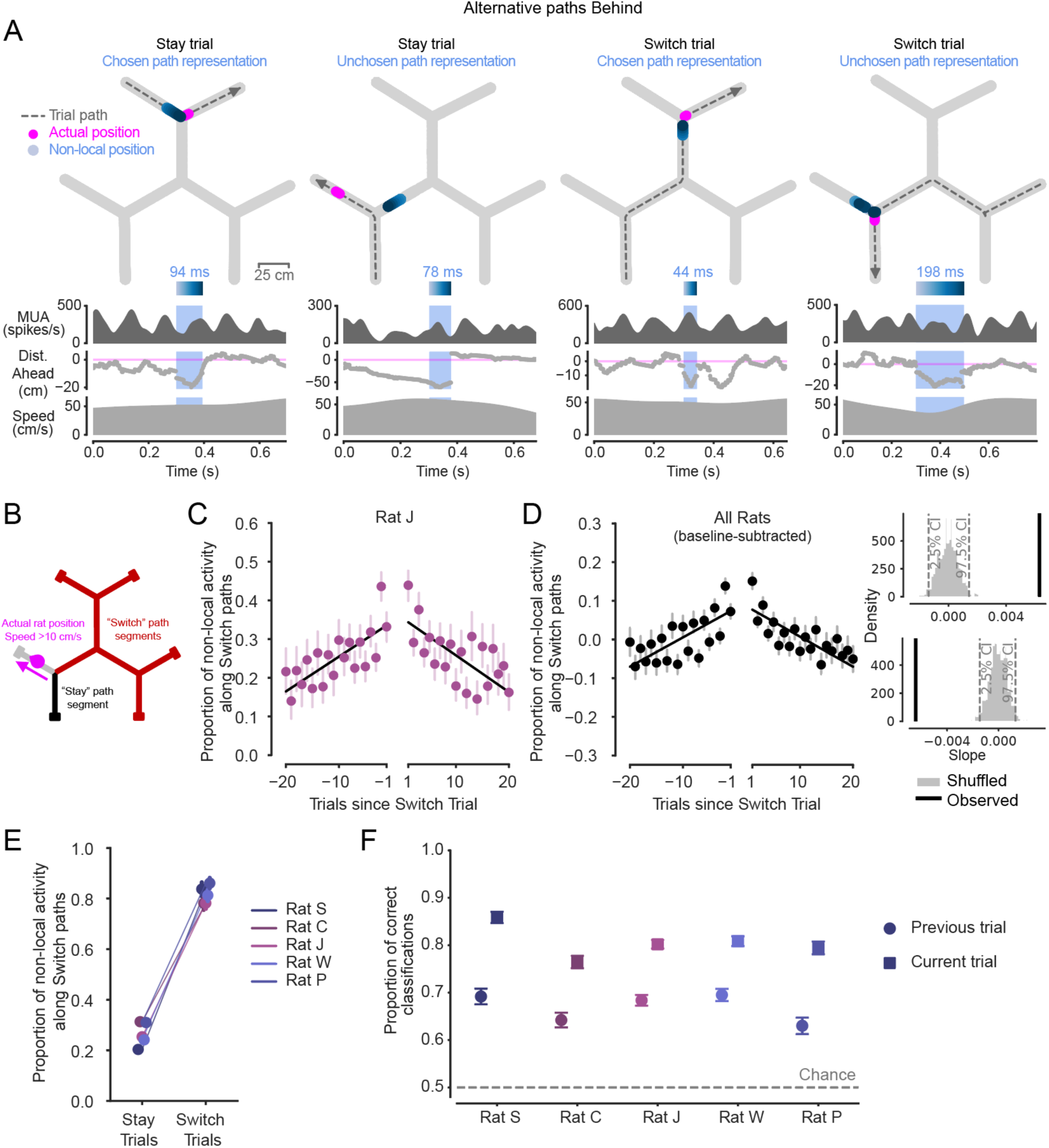
Non-local representations of alternative paths Behind are also enriched across trials before and after patch Switching. **(A)** Examples of non-local representations of paths behind in Stay and Switch trials. Top: Animal’s actual position (magenta) and decoded position from hippocampal spiking (blue, shaded by time) during the blue-highlighted period (below) are plotted on the track. Dashed grey line with arrow indicates animal’s path and direction on the current trial. Highlighted non-local duration is labeled above blue time-shaded heatmap. Top-middle: multiunit spike rate from hippocampal tetrodes before, during, and after highlighted non-local representation. Bottom-middle: Distance from actual position to decoded position, with positive values corresponding to in front of the animal’s heading, and negative numbers corresponding to behind the animal’s heading. Bottom: Animal speed. Note that periods outside the blue shaded region where the distance behind is large (e.g., second example) correspond to representations that are in the same track segment as the animal at that time rather than an alternative non-local segment. **(B)** Schematic defining Stay and Switch non-local activity behind, analyzed in **C-F**. Here, the relevant choice point is behind the animal, at the intersection between the grey, black, and red track segments. **(C)** Proportion of all non-local activity that represents paths consistent with Switching during the approach of the reward port across Stay trials preceding and following Switch trials in one rat. Left and right x axes separated to reflect patch change. Error bars are 95% CIs on the mean. Linear regression lines show increasing and decreasing proportions before and after patch Switch trials occur. Individual data for all rats shown in Fig. S2B. **(D)** Left: Proportion of all non-local activity that represents paths consistent with Switching during the approach of the reward port across all animals (n=5 rats), normalized per animal by subtracting off the average baseline proportion on Stay trials. Error bars are 95% CIs on the mean. Pre- and post-Switch linear regressions overlaid. Upper right: observed slope of pre-Switch regression (black) is significantly different from 0 (p<0.002) based on 1000 shuffles of the underlying data (grey). Lower right: observed slope of post-Switch regression (black) is significantly different from 0 (p<0.002) based on 1000 shuffles of the underlying data (grey). Both slopes were individually significant in all five animals (Fig. S2B). **(E)** Proportion of all non-local activity that represents paths consistent with Switching during the approach of the reward port in each animal on Stay trials and Switch trials. Error bars are 95% CIs on the mean. Switch trial distributions are significantly greater than in Stay trials for all animals (p=5.6e-264, 1.1e-155, 3.2e-300, 1.0e-100, 1.1e-277, Wilcoxon rank sum test, for n=292, 344, 525, 424, 443 Switch trials and n=3079, 3660, 6582, 4924, 3001 Stay trials). **(F)** Accuracy of logistic regressions predicting Stay or Switch choices based on proportion of all non-local activity that represents paths consistent with Switching during the approach of the reward port. Neural data from two trial types: previous trial (circle marker), or current trial (square marker). Error bars are 95% CIs on the proportion. Training data were balanced, so chance level was 0.5. Accuracy of model using previous trial neural data is significantly greater than chance in all animals (p=2.4e-58, 7.1e-32, 1.2e-56, 1.3e-40, 3.1e-48, Z test for proportions), and current trial model accuracy is greater than previous trial model accuracy (p=2.4e-58, 7.1e-32, 1.2e-56, 1.3e-40, 3.1e-48 ).

These non-local representations of Stay or Switch paths behind the animal (Fig. 3C,D) showed a very similar pattern of modulation around Switch choices as did non-local representations of paths ahead of the animal (Fig. 2C,D). Even though the representations behind were expressed after the animal had behaviorally indicated a choice on the current trial (Fig. 3B), the relative representation of the counterfactual Switch path ramped up across Stay trials before a Switch and ramped down across the subsequent Stay trials after arriving in a different patch (Fig. 3B-D, Fig. S2B). Again, this increasing and decreasing pattern was roughly symmetric on average around the Switch trial. Furthermore, just as for representations ahead (Fig. 2E), representations behind the animal were biased on average to represent locations along the actual path taken on the current Stay or Switch trial (Fig. 3E). Thus, the non-local representations behind the animal were systematically modulated with the evolving relative values of the options (Fig. 1F).

As with non-local representations extending ahead of the animal, the dynamic generation of representations of alternatives behind the animal was primarily driven by changing levels of Switch path representation (Fig. S3B). This is consistent with an increasing relative representation of the unchosen alternative path leading up to and following a choice to Switch. These results were also consistent when analyses were limited to representations extending at least 10 cm behind the animal (Fig. S3D), as well as in bouts within a patch lasting at least 10 Stay trials (Fig. S3G). Further, these non-local representations behind were concentrated in early phases of the theta rhythm, as expected from previous work^28,29,37,58^ (Fig. S2D). As for representations ahead, these results also could not be explained by spiking rates and were consistent when analyses were limited to continuous-state decoding periods (see Methods).

Representations behind the animal could also predict future behavior, consistent with an across-trial decision process. The relative representation of non-local paths behind the animal on the previous trial were predictive of whether the animal chose to Stay or Switch on the entirely subsequent trial (Fig. 3F). Here, the model predictors correspond to the proportion of non-local representation corresponding to Switch versus Stay paths, which is the same metric as in Figs. 3C-E. These Stay and Switch choice predictions were, as expected (Fig. 3E), not as accurate as those from non-local representations occurring as the animal traversed the final segments of trials, after the animal had already behaviorally expressed a choice to Stay or Switch (Fig. 3F). Nonetheless, previous trial non-local representations predicted subsequent choice well above chance. Thus, the proportion of non-local representation of the Switch path behind the animal on a given trial could predict what the animal would do several seconds later following traversal down the track segment to the reward port and back up the track segment to the choice point.

Together with the results from representations ahead of the animal (Fig. 2), these findings indicate that animals can progressively engage non-local representations associated with a progressively more valuable unchosen alternative (here Switching) both before and after the choice point leading to that option on each trial. These findings provide further support that the hippocampus dynamically samples alternative options, and can do so across successive trials with varying cognitive demands throughout flexible decision making.

### Representations of remote alternatives

Beyond retrieving information from an internal model related to an ongoing decision-making process, non-local information could be useful for additional purposes, including relating experiences or making inferences across space. In this task, where reward ports are distributed far across the maze, representing distant locations^38,43,61–63^, including those in alternative patches, could support relating reward experiences across space or updating relative values of locations based on local reward experiences. We therefore asked whether there was evidence for representations of distant locations across the maze, and whether these representations were specifically generated in relation to patch Switching, as animals learned from dynamic reward experiences.

In addition to non-local representations corresponding to track segments neighboring the animal’s actual location (Fig. 2A,3A; Fig. 4A left), we also observed non-local representations corresponding to alternative patches, closer to other reward ports (Fig. 4A, Supplemental Video 3). Alternative patches were represented on 10-20% of trials with any non-local representation (Fig. S4C). To quantify the extent of non-local representations, we then measured the maximal distance in centimeters between the animal’s actual position and the most likely decoded position represented by the hippocampus on each trial, during running at least 10 cm/s (Fig. 4B). Given our previous observation of representations of paths both ahead (Fig. 2) and behind (Fig. 3), we investigated non-local distance both as animals approached the first choice point and after the animal passed the final choice point in each trial (Fig. 4B).

**Figure 4.**
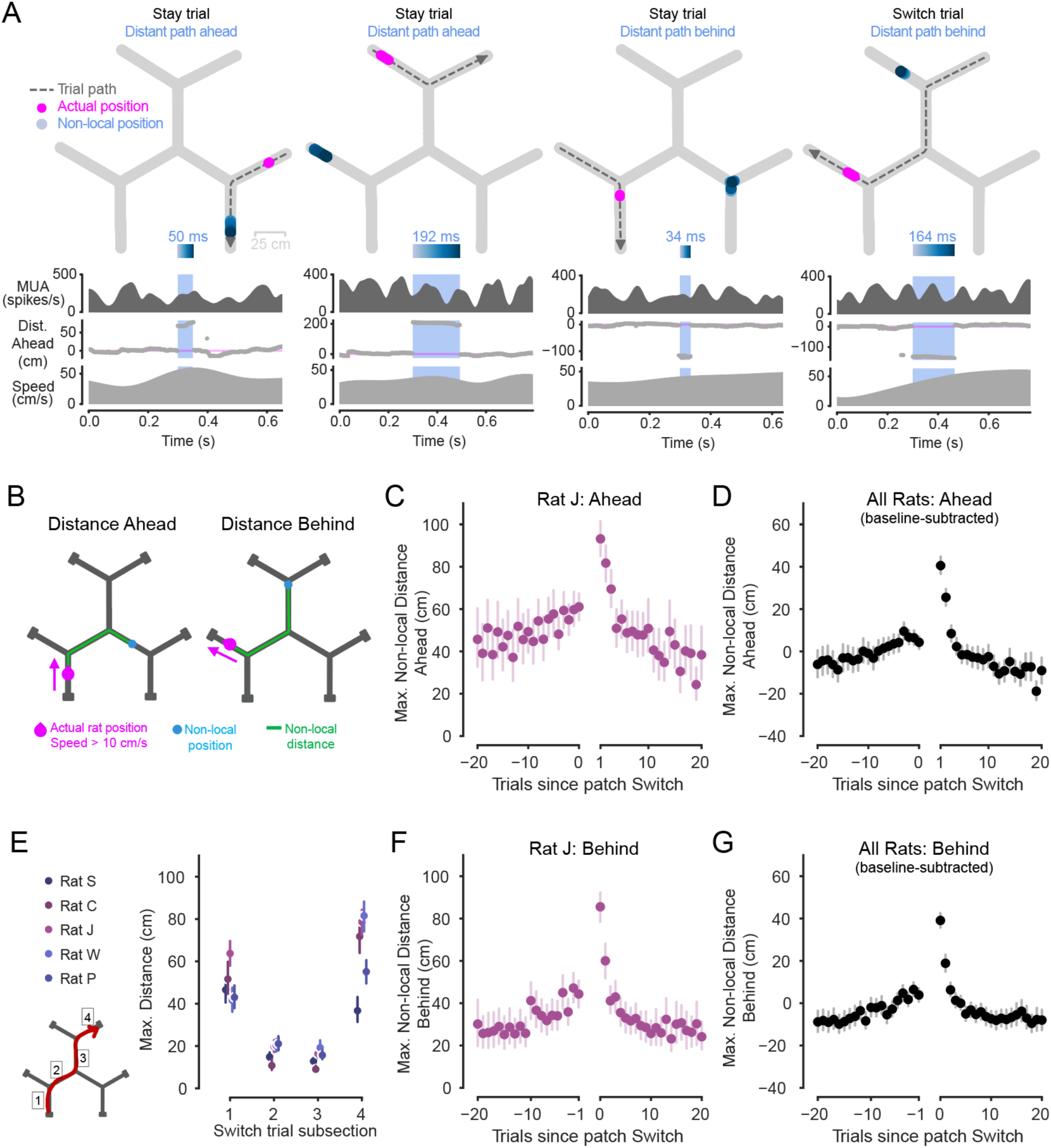
Non-local representations extend further early in patch experience. **(A)** Examples of distant non-local representations of paths ahead or behind, in track segments distinct from the animal’s current segment. Top: Animal’s actual position (magenta) and decoded position from hippocampal spiking (blue, shaded by time) during the blue-highlighted period (below) plotted on the track. Dashed grey line with arrow indicates animal’s path and direction on the current trial. Highlighted non-local duration is labeled above blue time-shaded heatmap. Top-middle: multiunit spike rate from hippocampal tetrodes before, during, and after highlighted non-local representation. Bottom-middle: Distance from actual position to decoded position, with positive values corresponding to in front of the animal’s heading, and negative numbers corresponding to behind the animal’s heading. Bottom: Animal speed. Representations can correspond to distant locations within the current patch (left example) as well as in alternative, unoccupied patches (right three examples). **(B)** Schematic showing non-local distance, defined as distance in centimeters of the path along the track (green) from the animal’s actual position (magenta) to the peak of the decoded posterior (blue). Example schematics correspond to representations of locations an example distance ahead of (left) or behind (right) the animal’s actual position. **(C)** Maximum non-local distances represented on trials leading up to and following patch Switches for one animal. Data taken from during the period of each trial in which the animal approached the first choice point (as in **B**, left). Left and right x axes separated to reflect patch change. Error bars are 95% CIs on the mean. Individual animals shown in Fig. S4A. Exponential regression intercepts *a* and rate constants *b* are significantly larger magnitude for post-patch-change than pre-patch-change data in all individual animals except one (p*_a_* = 1.3e-4, 4.6e-9, 1.1e-11, 3.1e-6, 0.8, p*_b_* = 2.3e-4, 1.1e-8, 3.2e-10, 3.3e-6, 0.9, Z test). Rate constants per animal indicated increasing and decreasing patterns before and after patch changes, respectively, in all animals (*b*_pre_ = 0.0172, 0.0144, 0.0191, 0.0105, 0.0185, *b*_post_ = -0.0177, -0.0388, - 0.0486, -0.0421, -0.0449, all p<0.05). **(D)** Maximum non-local distance ahead represented on trials leading up to and following patch Switches, across all animals. Data taken from first segment of trials and normalized per animal by subtracting off the average across trials. Error bars are 95% CIs on the mean. **(E)** Maximum non-local distance represented for each animal while traversing each of the four segments of a Switch trial (segments 1-4 schematized in lower left). Error bars are 95% CIs on the mean. Non-local distance is significantly greater when each animal is in the first segment than the second (p=4.7e-52, 1.0e-55, 6.4e-59, 8.5e-24, 2.1e-42, Wilcoxon rank sum test), and is also significantly greater in the final segment than the third segment for each animal (p=2.3e-49, 2.4e-56, 3.3e-69, 1.5e-49, 3.0e-70). **_(F)_** Maximum non-local distance represented on trials leading up to and following patch Switches for one animal. Data taken from final segment of each trial, during approach of the reward port (as in **B**, right). Left and right x axes separated to reflect patch change. Error bars show 95% CIs on the mean. Individual animals shown in Fig. S4B. Exponential regression intercepts *a* and rate constants *b* are significantly larger magnitude for post-patch-change than pre-patch-change data in all five animals individually (p*_a_* =8.8e-9, 2.9e-9, 1.4e-7, 3.2e-7, 9.2e-4, p*_b_* =5.7e-6, 3.0e-7, 2.3e-5, 4.0e-5, 3.6e-2, Z test). Rate constants per animal indicate increasing and decreasing patterns before and after patch changes, respectively, in all animals (*b*_pre_ = 0.0106, 0.0108, 0.0133, 0.0115, 0.024, *b*_post_ = -0.0206, -0.0384, -0.0344, -0.0336, - 0.0478, all p<0.05). **(G)** Maximum non-local distance behind represented on trials leading up to and following patch Switches, across all animals. Data taken from final segment of trials and normalized per animal by subtracting off the average across trials. Error bars are 95% CIs on the mean.

We found that non-local distance ahead was strongly modulated (Fig. 4C,D) in a pattern surprisingly distinct from the modulation of non-local content (Fig. 2C,D). The maximum non-local distance gradually ramped up across Stay trials and the first segment of Switch trials (Fig. 4C,D). The extent was then greatly elevated once the animal arrived in a new patch, and then decreased rapidly across the subsequent few Stay trials (Fig. 4C,D). This is in marked contrast to the modulation of non-local content, which increased and decreased in a roughly symmetrical manner. The maximum non-local distances could reach an average of approximately 90 cm from the animal (Fig. 4C, Fig. S4A), and, across animals, increased by approximately 40 cm from baseline (Fig. 4D) on the Stay trial following a Switch. This asymmetric pattern of non-local extent around Switch trials suggests a second kind of modulation of the generation of non-local representations across successive trials by the hippocampus during experience-guided decision making.

We then examined how these distances changed throughout the Switch trials themselves (Fig. 4E). Here we considered maximally distant representations both ahead and behind the animal during traversal of each of the four track segments of Switch trials. We found that while on average non-local representations extended as far as 40-60 cm from the animal on the first segment, this distance dropped to approximately 20 cm on both the second and third segment. Then, the maximum distances rose again to approximately 40-80 cm as the animal traversed the final track segment of the trial and approached the reward port in the new patch. This is consistent with the geometry of the environment, with the available space ahead or behind the animal changing as the animal progresses through the track segments, and adds to previous work indicating that hippocampal activity can be modulated by many additional factors including the distance to a goal^43^ or choice point^38^, rewarded or salient locations^64–73^, and familiarity versus novelty^8,74,75^.

The extent of non-local representations behind the animal also showed an asymmetric pattern of across-trial modulation around Switches (Fig. 4F,G, Fig. S4B). That is, the maximum distance of non-local representations behind the animal on the final segment of trials was also elevated upon Switching patches and then decreased back to baseline over subsequent Stay trials (Fig. 4F,G). Thus, the extent both ahead and behind showed a modulation pattern around Switch trials that was distinct from the modulation of Switch versus Stay path content (Fig. 2,3). Here again spiking rates could not explain our findings, and all results were consistent when analyses were limited to continuous-state decoding periods (see Methods).

We next asked whether non-local representations that corresponded to alternative, unoccupied patches were predictive of past or future patch choices. We found no evidence for such a relationship. Representations of specific alternative patches were neither biased to represent the patch the animal just arrived from nor predictive of the patch the animal would visit next (Fig. S4D). And, while in some animals these alternative patch representations did tend to overrepresent the track segments containing reward ports, rather than the central three track segments, this effect was not consistently observed across animals (Fig. S4E). These results indicate that representations of specific distant spatial alternatives in other patches are not directly reflective of specific prior or upcoming Switch choices.

Together, these patterns indicate that non-local extent can be specifically modulated across trials in patterns distinct to those observed for non-local content. These findings also raise the possibility that more distant representations are involved in computations particularly required upon patch switching, including the initial approach of the first reward port in the new patch. This is a period in which the animal can learn about the reward values in the new patch, and potentially how they relate to values across the maze.

### Non-local representations extend during learning opportunities

The observation that the extent of non-local representations was greatly elevated on the final segment of a Switch trial and for a few trials thereafter, after entering the new patch, suggested that representing distant locations might be particularly useful at those times. We reasoned that, as animals made their Stay and Switch decisions based on both local and remote patch values (Fig. 1E), a mechanism that enabled the retrieval of the value of alternative remote locations would be beneficial. Specifically, if, after entering a new patch, the animal could access the value of remote locations, it could then compare the current estimated value of Staying to that of Switching. Moreover, if the animal’s estimated value of Staying in a patch were uncertain and changing rapidly based on new reward experiences, then updated comparisons could be especially informative for making adaptive decisions moving forward.

To investigate how non-local distance varied at the level of single trials with any patterns in these model-inferred animal beliefs about port values, we next combined the data for representations ahead and behind the animal (Fig. 4D,G) into a single value of the non-local distance for each trial. As expected, the resulting average distances showed a large increase on the Switch trial, a peak on the following trial, and then a rapid decline over subsequent trials (Fig. 5A).

**Figure 5.**
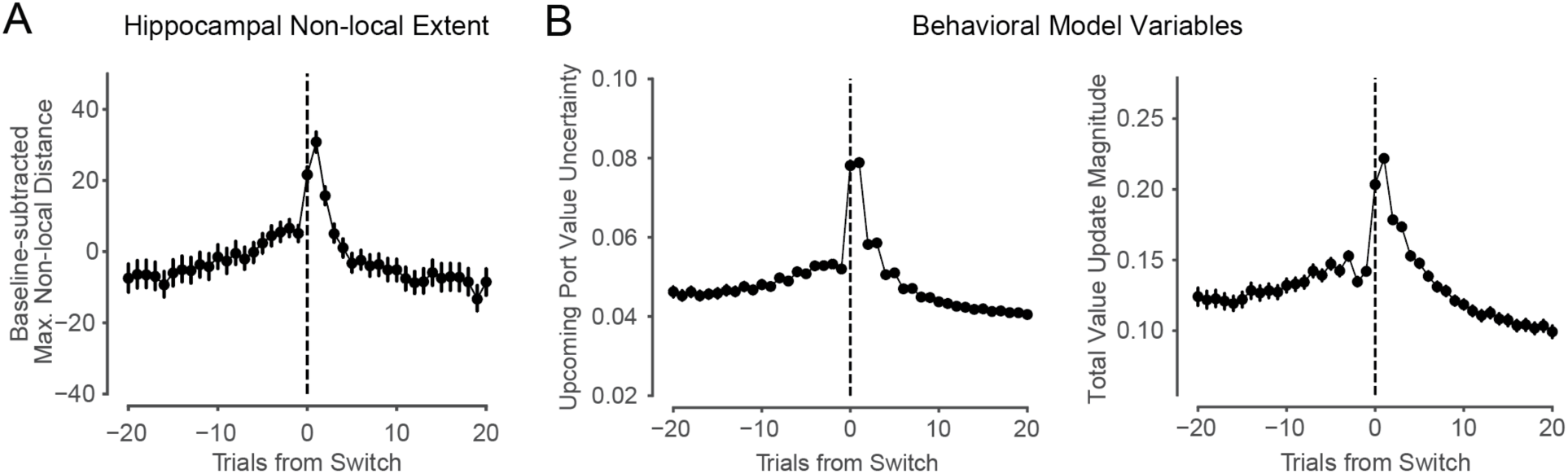
Behavioral modeling captures enhanced learning opportunities in early patch experience. **(A)** All animals’ normalized maximum hippocampal non-local distance represented on trials leading up to and following Switch trials, for non-local representations both ahead and behind (as in Fig. 4D,G). Trial-level quantification, rather than sub-trial-level quantification, shown here to match trial-level resolution of behavioral model. Error bars are 95% CIs on the mean. **(B)** Behavioral model variables related to value learning opportunities are enhanced upon patch Switching. Left: Uncertainty about the value of the upcoming reward port, measured by the value distribution’s variance, shows asymmetrical pattern around Switch trials, becoming elevated on and after Switch trials, and decreasing across trials thereafter. Patterns were consistent across animals, shown individually in Fig. S5F. Right: Absolute magnitude of value updating after each trial, measured as the change in value distribution means. Degree of value updating is elevated upon patch Switching, and decreases across trials the longer the animal Stays within a patch. Patterns were consistent across animals, shown individually in Fig. S5G. Error bars are 95% CIs on the mean.

In this scenario, we would expect a preponderance of distant representations upon entry into a patch (Fig. 5A) to occur during the period where the value of the newly entered patch was being updated rapidly. To test this prediction, we returned to the behavioral model. As mentioned above, the model inferred from animal behavior a probability distribution of reward likelihood at each port based on the animal’s previous reward experiences at that port. These beliefs about the chosen port’s value were updated trial-by-trial according to reward outcomes (Fig. S5A-C). In parallel, the value estimates for unvisited ports became less certain over time, decaying toward a uniform distribution. This captures the intuition that the certainty of beliefs about the values of locations that have not been visited for some trials would decrease over time. Thus, when the animal chose to Switch patches, it may visit new ports of uncertain value, and observing new reward information could have the potential to drive large updates to the animal’s beliefs about the value of the visited location. We therefore measured both value uncertainty and the degree of value updating across trials around Switch choices.

We observed that the model-inferred value uncertainty at the upcoming port peaked on the Switch trial and the subsequent trial and decreased over trials thereafter (Fig. 5B left). Moreover, the total estimated value update occurring on each trial showed a pattern of change similar to that seen in the maximal distance of non-local hippocampal representation: it rose substantially on the switch trial, increased further on the next trial, and then decayed back to baseline levels (Fig. 5B right). These patterns were consistent across all animals (Fig. S5F,G). As expected, value updates were predominantly driven by value learning at the visited local port (Fig. S5H). Taken together, these results are consistent with the idea that the hippocampus preferentially samples more distant locations across the environment during trials when expected values are uncertain and reward outcomes can guide learning about local value. This suggests that non-local distance will be enhanced during periods in which new reward outcome information must be integrated into the animal’s internal model of the environment.

## Discussion

Our findings demonstrate the distinct modulation of the content and spatial extent of non-local representations of alternatives in the hippocampus, in patterns that are well-matched to the animals’ evolving cognitive needs throughout flexible decision making. The task structure encouraged animals to continually update their estimates of the values of different options based on experience and to use those values to decide whether to Stay in the current patch or Switch to a different patch. Moment-by-moment decoding of hippocampal spatial representations during navigation, in combination with behavioral modeling, revealed two distinct patterns of modulation: (1) non-local representations of alternatives were expressed in relation to their relative expected values, with the higher value alternative making up the greater fraction, and (2) non-local representations extended further from the animal at times where our model suggested that value learning was most prevalent. These patterns of modulation spanned successive trials and engaged non-local representations of locations not only ahead of, but also behind the animal. Further, the representations included locations that were not only close to, but also very distant from the animal. These findings reveal the broad representational capacity of the hippocampus during locomotion and indicate that this representational capacity is specifically engaged in patterns appropriate to subserve cognitive demands associated with both making decisions among and updating internal representations of alternatives.

### Modulation of non-local representations with relative choice value

Previous work on non-local representations and cognitive functions during locomotion has often focused on representations ahead of the animal and identified a variety of relationships between the alternatives represented and current trial behavior. These results included an association between alternative representation content and head-scanning behavior^44^, although non-local representations are also observed in the absence of this behavioral signature^35,76^. These results also included reports that representations of specific alternatives either did^41,43,45^ or did not^35,40,42^ predictably relate to the upcoming choice on the current trial.

Our results provide a potentially unifying framework to explain these previous observations. We identified a modulation of non-local representational content with relative value across successive trials. Specifically, when the difference between the values of the Stay and Switch options was large, the higher value alternative dominated non-local representations. When the differences were smaller, the relative proportion of non-local representation of each was more similar. These findings suggest that the expected values of options or related decision variables influence the extent to which those options are expressed in the context of non-local activity.

This provides a potential prioritization of options to be considered, as the hippocampus would be most likely to represent the best, most viable options. Similar prioritization, in which the most promising options are internally sampled most extensively, also arises in rational computational analyses of choice under uncertainty^60^.

Thus, when the preferred choice is clear, the hippocampus primarily represents that path, but nevertheless samples occasionally from the alternative, potentially to retrieve information about the path not taken. In conditions where the preferred choice becomes less clear, the hippocampus generates progressively more balanced representations of the two options. Critically, therefore, hippocampal non-local representations are neither constantly predicting the planned choice, nor strictly providing a menu that evenly represents all potential options— rather, representations can be more or less predictive of behavior depending on the current decision-making demands.

At the same time, relative value-related modulation of non-local representations suggests that the mechanisms selecting non-local representations already have an estimated value of the alternatives. If so, then why would it be useful to generate a non-local representation? We suggest that non-local representations activate place and associated value representations on the actual path toward possible goals. This sampling of place and value could progressively refine and provide a more accurate estimate of the value, and thus serve to better inform decision processes^10,46^. This is consistent with our observation that the proportion of representation of the Switch path ahead was predictive of choice on even the next trial, and that similar patterns of non-local content extending behind the animal also related to upcoming choice, well in advance of physically approaching the next choice point.

### Distinct modulation of non-local content and extent

We identified a surprising dissociation between the representational content and the spatial extent of non-local events, where representations of nearby portions of the un-chosen Switch path increased before a Switch decision, while representations of distant locations increased after the Switch decision. Our behavioral modeling framework provides a potential interpretation of these results.

In the trials leading up to a patch Switch, hippocampal representations content most often corresponded to locations along adjacent track segments rather than at remote port locations. This is consistent with a key idea of many reinforcement learning models—that choices are guided by a learned representation of the long-term consequences of some local action. In this framework, an animal would evaluate Stay or Switch options leading up to a Switch choice by retrieving the up-to-date values associated with the neighboring paths nearby rather than their ultimate remote destinations.

Our results also suggest that non-local representations of distant locations may particularly be recruited when retrieval is required for updating, potentially to support an internal computation of relative value. Behavior in the task depended on both local and non-local patch values, suggesting that the animals were retrieving and comparing these values to inform their decisions. Retrieving these values to enable an up-to-date comparison might then be particularly important when local values were uncertain and changing substantially due to new reward outcomes, which occurred upon entering a new patch and a few trials thereafter. Consistent with those possibilities, representations of distant locations were much more prevalent on and after Switch trials and showed an overall pattern of expression that closely matched behavioral model estimates of local value updating. One possibility is that these events enable caching of the estimated values of distant locations to inform Stay and Switch decisions. Additionally, it is possible that accessing information about remote locations supports a form of trial-by-trial updating that propagates local value information or relative value information across the environment, which could also then be accessed in subsequent decisions. Recordings and analyses focused on value representations across the brain will be required to assess this possibility.

While the potential mechanisms by which non-local representations during navigation could support retrieval and use of value information merit further investigation, some initial points can be made. First, the extent of non-local representations was low as animals traversed the central segments on a Switch trial but increased substantially in the final segment; that is, non-local extent increased immediately before rather than only after the first outcome in the new patch was revealed. This suggests that the role of these remote representations is not dependent on having just experienced an outcome at a reward port. Instead, the hippocampus may facilitate post-outcome decision-making or updating by retrieving representations of distant alternatives that are later used for a comparison. Alternatively, prior work has demonstrated that non-local representations during navigation are strongly associated with the theta rhythm, and sequences along the theta rhythm are thought to support learning by binding together elements of experience at a compressed timescale through plasticity mechanisms^38,48,58,77–79^. Thus, this system could be used to learn relationships between current experience and unchosen alternatives within theta cycles (or other organizing codes^80^), in support of updating knowledge of remote alternatives during active navigation. This hippocampal work leaves open the question of how non-local spatial representations may be coordinated with value-related learning and decision processes across the brain ^6,81–85^.

### A broad range of alternatives are flexibly engaged

Our results also establish that a diverse range of alternatives are expressed by hippocampal non-local representations during active navigation. In the context of a specific decision among options, previous work has often focused on representations that corresponded to possible future locations on the current trial^40,42,43^. Our findings show that representations of alternative paths not only ahead of the animal but also behind the animal—even when moving away from the associated choice point—are flexibly engaged across trials associated with decision-making and learning processes. We also found that this engagement included non-local representations of very remote locations, including representations that occasionally jump far across the maze rather than sweep ahead at a constant rate.

The dynamic engagement of a broad range of alternative possibilities by the hippocampus^35^ during locomotion is consistent with the idea that non-local population representations can support a range of cognitive functions that involve computations about alternatives^86–89^. In particular, the dynamic engagement of representations behind the animal^38^ during navigation raises the possibility that the hippocampus supports computations about actual and counterfactual^90^ past paths during navigation, not only during rest. Moreover, the relative prevalence of representations behind the animal predicted behavior on the subsequent trial, indicating that representations of alternatives behind the animal need not be strictly retrospective, but could also support future decision making.

Additional observations from prior studies exemplify that hippocampal activity during locomotion can correspond to various correlates, including representations of opposite directions^35^, forward and reverse sequences^91^, distant locations^43,61^, alternative environmental contexts^62^, and different spatial reference frames^63^; our results are consistent with the idea that any of these representations of alternatives may be differentially engaged and specifically curated across trials depending on the cognitive computations needed for the task at hand.

### Non-local computations in immediate and long-term adaptive behavior

Retrieving information about alternative options related to making a specific upcoming decision and retrieving information about alternatives related to updating an internal model are both computations that inform inference across space (“non-local inference”). In the former case, representations of alternatives may support non-local inference of which nearby alternative path has the greater expected value based on prior experience. And, in the latter, representations of alternatives may support non-local inference about the greater remote or local value, or how the expected relative value of locations should be altered based on outcomes observed locally. This possibility is consistent with prior work indicating that the hippocampus is important for inference^7,92–100^, and relational thinking more broadly^26,101–105^.

The prolonged relationship across trials of non-local representations to Switching behavior serves as a reminder that the behavioral benefit of non-local inference need not be immediate. While we observed that non-local representations can predict immediate future and past Stay or Switch choices, we also found that representations of alternatives ramped across many trials before and after Switch choices, that they predicted choices an entire trial in advance, and that remote representations in alternative patches did not predict prior or subsequent patch choices. Thus, while non-local representations can have immediate behavioral benefit by supporting the immediately upcoming choice, they may also support decision making over the longer term, perhaps by integrating and accumulating^106,107^ retrieved information across trials. This is consistent with the idea that inferring structure in an environment and relations between experiences may occur in the background of behavior^108^, particularly as new information needs to be incorporated into an internal model, even if an animal does not yet know if or when it will later need to draw on that knowledge to support adaptive choices. This is thought to be especially important in complex and dynamic environments to enable flexible decision making when confronted with new or unexpected circumstances.

While the circuit-level mechanisms that determine the flexible generation of representations of alternatives remain largely unstudied, our observations of modulation with respect to relative values and the need to learn them implicates brain regions like the prefrontal cortex as possible drivers^81–85,109–112^. Consistent with this possibility, activity in the medial prefrontal cortex (mPFC) and the hippocampus can be precisely coordinated^42,113–115^ and mPFC spiking can predict the future engagement of non-local hippocampal spiking^61^. Further, observations of coordination of hippocampal representations and peripheral sensory-motor processes^116^ suggest broad engagement of many brain areas at times when hippocampal non-local activity may be important for ongoing behavioral computations. The specific generation of non-local hippocampal activity patterns and associated representations of alternative possibilities could engage widely distributed computations to enable querying internal models for learning and adaptive decision making in a complex and dynamic world.

## Resource Availability

### Lead Contact

Requests for further information should be directed to the lead contact, Loren Frank.

### Materials Availability

This study did not generate new unique material reagents.

### Data and Code Availability

Data will be available on the Distributed Archives for Neurophysiology Data Integration (DANDI) Archive at https://dandiarchive.org (ID 001942). An interactive version of the paper will be available at https://doi.org/10.62329/yhwc6622. Code will be available at https://github.com/LorenFrankLab/comrie_et_al_2024. Additional information is available upon request from the lead contact.

## Acknowledgements

We thank Anya Kiseleva, Victor Perez, and Mushu Yusifova for administrative support, Viktor Kharazia for assistance with histology, Tom Davidson, Anna Gillespie, Hannah Joo, Clay Smyth, Maxim Borius, and Cameron Wilhite for early technical assistance, Shijie Gu for assistance with behavioral modeling, Samuel Bray and Chris Brozdowski for assistance with data sharing, and Kenneth Kay, Daniel Silversmith, Vanessa Bender, Alexandra Nelson, and Michael Brainard for guidance, discussions, and comments on previous versions of this work. This work was supported by NSF GRFP 1650113 (A.E.C.), NIH F31MH124366 (A.E.C.), UCSF Jonas Cohler Discovery Fellowship (A.E.C.), the Life Sciences Research Foundation (A.J.), NIH F30MH126483 (J.A.G.), NIH R01MH121093 (N.D.D.), NIH R01MH136875 (J.D.B., N.D.D., L.M.F.), Simons Collaboration on the Global Brain 521921 and 542981 (L.M.F.), and Howard Hughes Medical Institute (L.M.F.).

## Author contributions (CRediT taxonomy)

Conceptualization, A.E.C., L.M.F.; Methodology, A.E.C., A.E.K., T.A.K., J.D.B., N.D.D., L.M.F.; Software, A.E.C., E.J.M., A.E.K., E.L.D., N.D.D., L.M.F.; Formal Analysis, A.E.C., E.J.M., A.E.K.; Investigation, A.E.C., E.J.M., A.J., J.A.G.; Resources, L.M.F.; Data Curation, A.E.C., E.J.M., E.L.D., L.M.F.; Writing – Original Draft, A.E.C., L.M.F.; Writing – Review & Editing, A.E.C., E.J.M., A.E.K., E.L.D., A.J., J.A.G., T.A.K., J.D.B., N.D.D., L.M.F.; Visualization, A.E.C., A.E.K.; Supervision, L.M.F.

## Methods

### EXPERIMENTAL MODEL AND SUBJECT DETAILS

#### Subjects

All animal procedures were approved by the Institutional Animal Care and use Committee at University of California, San Francisco. Five adult male Long-Evans rats (Charles River Laboratories, 450-650g) were each pair housed in a temperature-and humidity-controlled environment on a 12-hour light-dark cycle with lights on 6 AM - 6 PM. Rats had *ad libitum* access to standard rat chow and water. Prior to behavioral training, rats were transitioned to single housing and food restricted to 85% of their baseline weight.

### METHOD DETAILS

#### Behavioral pre-training

All behavioral tasks were controlled via custom code written in Python and Statescript in combination with an Environmental Control Unit (Spikegadgets). Animals were pre-trained to run back and forth on a linear track with walls for liquid food reward (Carnation evaporated milk sweetened with 5% sucrose) from reward ports fixed at each end of the track. Upon entry of a reward port, 100 μL reward was delivered by syringe pump immediately and automatically, gated by an infrared beam. Animals learned to alternate between and obtain rewards from the two ports across two 40-minute and one 25-minute sessions. To familiarize animals with navigating an elevated maze, animals then performed the same task on an elevated linear track (1.1 m long, 84 cm high) until they performed at least 30 rewards in a 15-minute session. Pretraining took place in an environment with distal spatial cues on the walls. The pretraining environments were fully separate from the Spatial Bandit Task environment. Animals were subsequently returned to *ad libitum* food access prior to surgical implantation.

#### Neural implant

Custom hybrid microdrives contained 24 independently movable 12.5 μm diameter nichrome tetrodes (California Fine Wire and Kanthal). The drive body was 3D printed (PolyJet HD, Stratasys) and funneled tetrodes into two cannulae to target 12 tetrodes to each hemisphere. Tetrodes were gold plated to reduce impedance to ∼250 kOhms. Implants also contained a custom headstage (Spikegadgets) that coupled with a stacking set of up to four custom 128-channel polymer probes^117^ (Lawrence Livermore National Labs) per animal. Implant was housed in a custom 3D-printed enclosure with removable cap. Drive was sterilized before surgical implantation.

Custom hybrid microdrives were implanted stereotaxically under sterile conditions. In anesthetized animals, cannulae were implanted above dHC (+/-2.6-2.8 ML, -3.7-3.8 AP relative to skull Bregma) and polymer probes were implanted in mPFC and OFC. A stainless-steel ground screw was implanted over the cerebellum to serve as a global reference. Titanium screws were placed in the skull to help anchor the implant, which was secured with Metabond (Parkell, Inc) and dental cement (Henry Schein).

After surgery, tetrodes were advanced deeper into the brain daily over ∼3-4 weeks. One tetrode per hemisphere was advanced to corpus callosum as a local reference and all other tetrodes were advanced to dorsal hippocampus CA1 stratum pyramidale, guided by physiological signatures of spiking activity and the local field potential.

After full recovery from surgery, animals were food deprived to 85% of their baseline weight and reintroduced to the elevated linear track, as described above. Animals reached criterion again over several days to refamiliarize them with obtaining reward on a track and to familiarize them with running with their implant. Two rats had additional experience on a fork maze^118^. Re-training animals to run after surgery recovery ensured that the animals were motivated enough to begin the main experiment.

#### Data collection: Spatial Bandit task

The Spatial Bandit task took place on a maze with three radially arranged Y-shaped foraging “patches” each containing a central hallway that bifurcated into two hallways that each terminated in a photogated reward port, resulting in two ports per patch and six total reward ports. Each port had a separate automated pump, as described above. Hallways were 6.5 cm wide and linear track segments were each 53 cm long. All hallways intersected at 120 degrees, such that the track had three-fold symmetry. The track was made of black acrylic (TAP Plastics) with walls that were 3 cm high, enabling animals to see distal spatial cues.

The behavior took place in a dimly lit room (2.4 m by 2.9 m) with black distal spatial cues on the white walls and a plastic black curtain separating the maze from the experimenter, rig, and computer. A ceiling-mounted camera (Allied Vision) centered over the maze recorded animal behavior at 30 frames/second and was synchronized with all other data via Precision Time Protocol. Before behavior began each day, a ring of red and green LEDs was mounted atop the implant to enable online head position and direction tracking via SpikeGadgets Trodes software.

Animals began the task only once tetrodes had reached stratum pyramidale. Neural data was recorded from the animals’ first experience of the Spatial Bandit maze and environment. The general structure of each day of recording began by moving rats from their home cage into a rest box in the same room as the Spatial Bandit maze. Run sessions were interleaved with rest sessions throughout the day; rats typically completed 7 or 8 ∼20-minute run sessions per day, interleaved with rest sessions of at least 30 minutes each. Rest sessions helped to maintain stable motivation by providing breaks throughout each day. Each day also started and ended with a rest session. These behavioral data were collected over 8-17 days per animal.

During each run session, an animal was first placed in the center of the track facing the same wall each time, and then navigated freely in the environment, which was otherwise fully automated by Trodes (SpikeGadgets) software and custom behavioral control scripts written in Statescript (SpikeGadgets) and Python. At a high level, each individual run session contained multiple reward contingency blocks. Each contingency defined the reward probability p(R) assigned to each of the six reward ports. Contingency blocks covertly changed when an animal completed a certain number of trials, or hit a 20-minute time limit per contingency, whichever came first. Contingency changes almost always occurred based on the trial limit, very rarely changed based on the time limit, and did not depend on any other performance metric. A trial was defined as the period from exiting one photogated reward port to exiting a different port. Therefore, each trial contained both the run between two ports and an outcome (100 μL reward or omission) at the chosen port. Reward was only available probabilistically if an animal poked into a distinct port; consecutive pokes at the same port were never rewarded.

Each animal’s first run session began with reward probability p(R) of 1 at all reward ports for 100 trials, followed by an uncued contingency change to p(R) of 0.5 at all ports for the next 100 trials. This first session exposed the animal to the environment, encouraged the animal to visit all reward ports, and introduced probabilistic rewards. After this session, run sessions were each 180 trials long and contained three reward contingencies that each lasted 60 trials for four animals. One animal had 160 trial sessions each containing two contingencies of 80 trials.

From the second run session onward, each reward contingency defined a p(R) of 0.2, 0.5, and 0.8 per port, such that each contingency had a best patch on average. Both ports within a patch could have the same or different reward probabilities, but the best patch combination was 0.5 and 0.8 (not 0.8 and 0.8) across the two ports in a patch. By having ports of two different values within a patch, animals were encouraged to spend some trials not only at high-value but also at nearby lower-value locations, enabling us to sample neural data as animals visited ports of a full range of values. Contingencies were pseudorandomized to counterbalance which patch was best, whether the left or right port within each patch was best, and whether the best, medium, and worst patch followed a clockwise or counterclockwise order around the track. Contingency changes and reward port values were never cued, so that animals had to make navigational choices based on memory of their prior experiences across the maze.

Continuous neural data, as well as digital inputs and outputs associated with beam breaks and reward pumps, were recorded during each rest and run session at 30 kHz in four animals and 20 kHz in one animal using Trodes version 1.8.2 (SpikeGadgets).

#### Histology

At the conclusion of behavioral experiments, animals were anesthetized with isoflurane, and small electrolytic lesions were made. One day later, animals were anesthetized and transcardially perfused with 4% paraformaldehyde. Brains were fixed *in situ* overnight, after which tetrodes were retracted from the tissue and brains were extracted from the skull. Brain fixation continued for two days. After cryoprotection by 30% sucrose for 5-7 days, brains were blocked and sectioned with a cryostat into 50-100 μm sections. Nissl labeling enabled identification of tetrode tracks and localization of tetrode tips at the sites of electrolytic lesions (Fig. S1B).

### QUANTIFICATION AND STATISTICAL ANALYSIS

#### Data processing and analyses

All data processing and analyses were carried out using Python and Julia using Spyglass^119^.

#### Behavioral analysis

Trials were parsed based on behavioral Statescript log files (SpikeGadgets). A trial was defined as the final exit from one port to the final exit from another port, and therefore included both the run from one port to the next as well as the entry into and outcome at the chosen port. Stay trials were defined as starting and ending within the same track patch, while Switch trials were defined by starting and ending in distinct track patches. The nominal reward probabilities assigned to each port and reward delivery times were also tracked by parsing log files. The nominal best patch on each trial was defined as the patch with the highest average of the nominal reward probabilities of the two ports within the patch (Fig. 1D). These reward probabilities are referred to as nominal because they are assigned by the reward contingency. Importantly, an animal’s experienced reward values at each port at any time in behavior are related to but not necessarily identical to the location’s nominal reward probability, given the self-paced nature of the task and stochasticity of the rewards in the environment.

#### Behavioral model

To estimate the rats’ expected beliefs about the values of each of the six ports, we developed a Beta-Bernoulli model. This model estimates, for each port, the likelihood that the port has any possible reward probability value based on past rewards and failures at the port. Analogous to Q-learning, this model estimates the expected value of each port, and updates that value depending on experienced outcomes trial-by-trial. To extend this to track uncertainty in beliefs about port values, the model is Bayesian and estimates the probability of a port having any underlying success rate. Specifically, the model treats reward outcomes at each port as independent Bernoulli processes and maintains a Beta posterior distribution over the reward probability of each port, naturally handling the task’s binary outcomes. Each reward outcome updates the distribution corresponding to the chosen port on each trial.

The Beta-Bernoulli model tracks the value of port *i* with two variables, *α_i_* and *β_i_*, that correspond to the number of rewards and omissions – or successes and failures – at port *i*. The distribution that models the value of each port can also be re-parameterized with two different variables, via a simple mean and standard deviation, *μ_i_* and *σ_i_*, eliminating the need to track counts of past outcomes and allowing a more biologically plausible neural implementation. Means and variances of the posterior value distribution at each port also provide measures of expected value and uncertainty at each port on each trial.

The reward probability estimate, akin to a mean Q value, of any given port *i* is given by 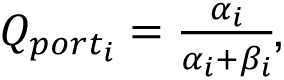 providing a simple estimate to compare port values.

The Q value of any given patch can also be computed from the values of the two ports it contains. The currently occupied patch Q value is computed as a weighted average of its two ports, with *Q_patc_*_ℎ_ = (1 − *γ*)*Q_upcomin_*_g_ + *γ* ∗ *Q_past_* where *Q_past_* denotes the Q value of the port the rat is departing from and *Q_upcomin_*_g_ denotes the Q value of the port the rat would next visit if it chose to Stay within the patch. This weighting allows for the possibility that animals considered the value of Staying in the current patch by taking the value of both its ports into account, rather than only considering the upcoming port value. For alternative patches, *Q_patc_*_ℎ_ was defined as the mean of the two port Q values within each patch.

### Behavioral model learning rule

Upon visiting a port, the model was updated according to the following rules.

After a reward at port *i*:

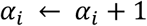

After an omission at port *i*:

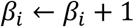

Both *α_i_* and *β_i_* corresponding to each port *i* were initialized to 1 at the beginning of each session.

To account for the possibility of forgetting when an animal did not visit a port for a while (Fig. S1A), both *α_i_* and *β_i_* of unchosen ports were slightly decayed towards 1 after each trial, slowly resulting in a forgetting of the unchosen port values, and a relaxation of beliefs toward a uniform distribution over possible reward probabilities.

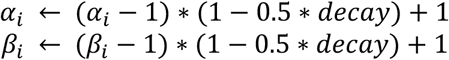

Note that while this learning process can be simply modeled through *α* (rewards) and *β* (omissions), it can also be easily reparameterized and equivalently expressed via *μ* (mean) and *σ* (standard deviation), which allows for a more straightforward neural implementation. Here the mean Q value of a port is:

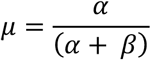

With variance:

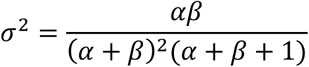

Importantly, this enables the learning rules themselves to be rewritten in terms of *μ* and *σ*. Specifically, the update rule to the mean can be written as a delta rule, akin to Q learning, with *r_t_* as the reward outcome observed on trial *t*:

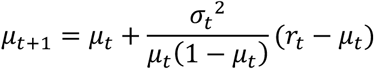

The variance is then scaled by a factor that depends on both the previous and new means:

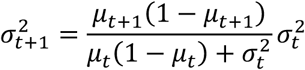

### Behavioral choice model

To connect this learning model to choices and assess our ability to capture animal behavior, we computed the likelihood of choice of patch on every trial as the softmax of the three patch values, given by:

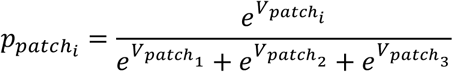

The value at the current patch was given by:

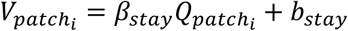

For the patch clockwise from the current patch, value was given by:

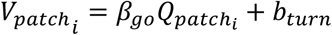

The value of the remaining patch was given by:

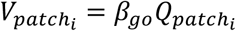

where *Q_patc_*_ℎ*i*_is the patch Q value estimated on each trial as described above.

The inverse temperature parameters, *β*_g*o*_ and *β_stay_*, determined how sensitive choice behavior was to differences in expected reward – high temperatures indicate high sensitivity to reward, while low temperatures indicate insensitivity. To allow for differential sensitivity to reward estimates at the current patch versus alternate patches, we fit two independent softmax temperatures, *β_stay_* for the current patch, and *β*_g*o*_ for the two alternative patches. This reflected tendencies to Stay in the current patch versus to Switch patches.

The term *b_stay_* reflected a fixed cost to changing patches due to the time and distance to reach a new patch and was thus added to the valuation of the current patch.

To incorporate tendencies of animals to prefer to turn in certain directions when changing patches, we included a turn bias *b_turn_*, an offset that was consistently applied to the patch to the left of the animal’s current position. If the rat was in patch 1, this was added to patch 2; if the rat was in patch 2, this was added to patch 3; if the rat was in patch 3, this was added to patch 1.

On trials where the animal switched patches, likelihood of port choice was modeled as a subsequent softmax between the two ports, with a separate softmax temperature *β_port_* applied to the per-port reward estimates:

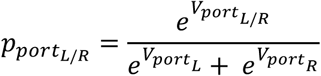

The value of the left port was given by:

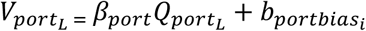

where *b_portbiasi_* was fit independently for each patch, reflecting turn biases in which reward port each rat tended to enter when switching to each patch.

The value of the right port was given by:

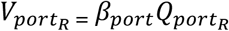

### Behavioral model fitting

For the Beta-Bernoulli model, we optimized the free parameters of the model (*γ*, *decay*, *β*_g*o*_, *β_stay_*, *b_stay_*, *b_turn_*, *β_port_*, *b_portbias_*) by embedding each of them within a hierarchical model to allow parameters to vary from day to day. These parameters were themselves modeled as arising from a population-level Gaussian distribution over days, separately per animal. In fitting the model, we added parameters one at a time and retained variables that improved the Akaike information criterion (AIC). We estimated the model for each animal to obtain best fitting day- and animal-level parameters to minimize the negative log likelihood of the data using an expectation-maximization algorithm with a Laplace approximation to the day-level marginal likelihoods in the M step^120^.

### Behavioral model variables

In subsequent analyses, we characterized value-guided Stay and Switch choices at the level of trials using variables estimated through the behavioral model, including the values associated with each port on each trial. We built a binomial family generalized linear model (Logit link function) for each animal to predict Stay or Switch choices from estimated expected values associated with the choice to Stay or to Switch (Fig. 1E). The value of Staying was defined as the model-estimated value of the upcoming port within the patch on the current trial, and the value of Switching was defined as the greater of the average port values in the two alternative unoccupied patches into which the animal had the option to Switch on the current trial. The difference between the Switch and Stay values on each trial, without accounting for the cost of Switching or other biases, defined the relative value of Switching and Staying on each trial (Fig. 1F, Fig. S5). The variance over the reward belief state of the upcoming port (Fig. 5B, Fig. S5) reflected a measure of uncertainty on each trial. The value update at each port on each trial was calculated as the difference between the port value before and after the outcome was observed on each trial, and absolute value updates were summed across ports where applicable to determine the total update magnitude on each trial (Fig. 5B, Fig. S5).

### Position

LEDs mounted on the front and back of the implant were tracked via overhead camera at 30 frames/second using Trodes software (SpikeGadgets). Positions were converted from pixels to centimeters, smoothed, interpolated and upsampled to 500 Hz to match neural decoding resolution. The front and back LEDs were used to calculate head orientation. Head angle was measured between the head orientation and the currently occupied linear track segment. The head position and head speed were calculated from the centroid of the font and back LEDs. To enable faster decoding, head position was linearized into one dimension by projecting the two-dimensional head position onto a graph representation of the track with nodes at the reward ports and track segment intersections, connected by edges. In between the track edges, which corresponded to the nine physical track segments, we introduced 15 cm gaps in 1D space to avoid inappropriate smoothing across adjacent linear positions that were not adjacent in physical space (Fig. S1).

### Spatial decoding

To decode spatial position from hippocampal spiking at high spatial and temporal resolution, we used a previously established clusterless state space algorithm^53,54,121–124,116^ from Denovellis et al., 2021^53^. Briefly, for each session, we built an encoding model to relate spike waveform features to the animal’s linearized head position in 2 ms bins using data from the beginning of the first trial to the end of the final trial of the session. The peak amplitude of each spike on each dCA1 tetrode channel served as spike waveform features. Spikes were detected by a 100 μV threshold on the recorded neural signal filtered 600-6000 Hz. To remove potential artifacts, a 1 ms window around times in which at least 75% of channels had an amplitude of 2000 μV or greater was excluded.

For each session, we then decoded spatial position from this hippocampal population spiking in 2 cm linear position bins and 2 ms temporal bins (Fig. S1). The state space decoding model^53^ had two states, spatially continuous and spatially fragmented, to capture smooth and discontinuous movement dynamics, respectively, of the hippocampal representation through the maze. The fragmented state was modelled by a uniform transition matrix and the continuous state was modelled by a random-walk transition matrix with a 6 cm movement variance. The probability of staying in the current state in the discrete transition matrix was 0.98. The initial conditions were set to be uniform across states and uniform across positions, as we did not have prior information about the most likely initial conditions. The model used kernel density estimators with a 24 μV bandwidth for waveform features and a 6 cm bandwidth for position to estimate the likelihood of a spike waveform feature predicting a position. Results were replicated after restricting data to periods of continuous-state decoding, with a threshold of at least 80% continuous state confidence, to rule out the possibility that fragmented states governed the reported patterns in the data. A similar decoding approach without position linearization was used for 2D spatial decoding for visualization purposes (Supplemental Videos 1-3).

The model output the posterior probability of position across the two movement dynamic states. We marginalized the joint probability over the two states, and then determined the most likely decoded position in each 2 ms bin as the peak of the posterior probability distribution.

We assessed decoding quality based on the concentration of the posterior probability, and only included times that were decoded with high confidence in subsequent analyses, by requiring the highest 50% of the posterior probability values to be concentrated within 50 cm of the track, as has been done previously^116^. We further note that this decoding approach was conservative in the sense that we decoded using the same data used to build the encoding model. In contrast, a cross-validated approach would instead decode periods distinct from those used to encode, and thereby exclude the place cell spikes predominantly associated with the animal’s actual current position during the decoded moments.

We measured the distance between the animal’s actual position to the most likely decoded position as the shortest path distance in linear space along the track in each 2 ms bin based on Dijkstra’s^125^ algorithm, with the animal and most likely decoded position as nodes on the track graph. We also classified the animal position and most likely decoded position into one of nine track graph segments in each 2 ms bin. For visualization, we projected linearized position back to the 2D track graph (Figs. 2A, 3A, 4A).

### Non-local representation of Stay and Switch paths

Non-local representation times were identified as times in which the most likely decoded position (hereafter “decoded position”) was in a track segment distinct to the animal’s actual position. We analyzed valid times of high speed (>10 cm/s) in order to limit our analyses to locomotor periods in which theta power is high and associated theta sequences are known to occur ^29,36,59,126^, and thereby excluded stationary rest periods in which sharp wave ripple replay events occur^1,8,34^.

To analyze the content of non-local representations occurring as the animal approached the first choice point on each trial (Fig. 2), we first identified which track segment the animal began in on each trial and used data from the time at which the animal exited the reward port to the first time at which the animal’s position exited that initial track segment. During this period, the animal approached the first choice point in the trial (Fig. 2B). Out of this period, we limited analysis to only the times that satisfied the decoding confidence and speed thresholds described above. For these valid times, we then quantified the total duration of non-local representation. We also quantified the duration of representation of the single segment ahead consistent with a subsequent Stay path and the duration of representation of any of the other segments ahead consistent with a subsequent Switch path, as in Fig. 2B. This enabled us to then calculate the proportion of the total valid non-local representation duration that was consistent with a non-local Switch path representation. The per-trial proportion was therefore calculated as: valid non-local Switch representation duration, divided by the sum of the valid non-local Switch representation duration and the valid non-local Stay representation duration.

A very similar approach was used to analyze the content of non-local representations occurring as the animal traversed the final track segment of each trial (Fig. 3). Here, we identified which track segment the animal ended in on each trial and used data from the final time at which the animal crossed from another segment into this final segment to the time at which the animal entered the chosen reward port. During this period, the animal approached the reward port, rather than a bifurcation (Fig. 3B). Again, we limited analysis to only the times that satisfied the decoding confidence and speed thresholds described above. For these valid times, we then quantified the total duration of non-local representation, and what proportion of that total non-local duration corresponded to a non-local representation consistent with a Switch path behind the animal (rather than the neighboring Stay path within the patch), as in Fig. 3B.

### Peri-Switch linear regressions and shuffles

Ordinary least squares linear regressions were fit to Stay trial data for trials before or after Switch trials (Fig. 2C,D, Fig. 3C,D, Fig. S2A,B). Shuffled distributions were obtained within animal by randomly shifting trial labels 1000 times, where the label is the number of trials from either the prior or next Switch trial.

### Predicting Stay and Switch choices from non-local representations

Generalized linear models with a logit link function (Fig. 2F, Fig. 3F) were used to predict Stay or Switch choices from the proportion of valid non-local representation corresponding to Switch paths. This proportion reflected the relative non-local representation of Switch versus Stay paths and is the same trial-by-trial metric as shown for representations ahead in Figs. 2C-E and representations behind in Figs. 3C-E. Models were fit separately per animal. Each model had one predictor, corresponding to the proportion of Switch path representation occurring while the animal occupied either the first or final segment of a trial t, the trial before t-1, or the average across the two trials t and t-1. Models were five-fold cross validated, and training data were balanced by resampling Switch trials. Predictions were binarized with a threshold of 0.5 and total accuracy was calculated across folds.

### Additional non-local representation generalized linear models

Generalized linear models were fit per animal using a binomial family with a logit link function. For Stay trials, the proportion of non-local activity corresponding to Switch versus Stay paths was modeled as a function of two predictors: the relative value of Switching versus Staying, to capture expected value information, and the number of trials away from the next Switch choice, to capture behavioral choice information even when the current trial choice was held constant. Relative value measures the behavioral model-inferred Switch value minus the behavioral model-inferred Stay value. In addition to quantifying model coefficients and significance, we also used the AIC to assess the contribution of individual predictors by comparing full models to reduced models where single predictors were excluded. For Switch trials, the proportion of non-local representation along Switch paths was modeled with relative Switch value as the only predictor.

### Theta phase

Neural data from a reference electrode in corpus callosum was filtered 0-400 Hz and downsampled to 1 kHz. This local field potential signal was then further filtered in the theta band from 5-11 Hz. The analytic signal was calculated with a Hilbert transform. The instantaneous phases for all valid local and non-local decoded run times were assessed separately for periods in which the animal was approaching the first choice point in each Stay or Switch trial, or after crossing the final choice point in each Stay or Switch trial (Fig. S2C,D). Phase convention refers to descending phases of corpus callosum theta as early phases, 0 to π, and ascending phases as late phases, π to 2π. Data were binned for visualization only. Full theta phase distributions for local and non-local times were compared with Kuiper’s non-parametric test for continuous circular data for each animal.

### Non-local distance

Non-local distance was calculated as the distance between the animal’s actual position and the most likely decoded position in each 2 ms bin (see spatial decoding methods). Using only high-quality valid running times, as described above, we identified the maximum absolute actual-to-decoded distance that occurred during the animal’s approach to the first choice point on a trial (Fig. 4C,D) or during the animal’s traversal of the final segment of a trial (Fig. 4F,G). Data in Fig. 4C,D,F,G were calculated based only on representations that occurred in non-local, unoccupied, track segments to limit representations to those either ahead of (Fig. 4C,D) or behind (Fig. 4F,G) the animal’s trajectory. To compare the non-local distance dynamics before and after Switches for each animal, we log-transformed the model *y* = *ae^bx^* into linear form and applied ordinary least squares regression to estimate log(*a*) and *b*, for both pre- and post-Switch data. Z tests were used to statistically compare the pre- and post-Switch parameters (Fig. 4C,D,F,G). Fig. 4E maximum distances were not required to be in a distinct non-local track segment to the animal’s actual position. Data occurring while the animal occupied the middle two segments of Switch trials (Fig. 4E) were calculated based on Switch trials in which exactly four segments were occupied. Wilcoxon rank sum tests were used for each animal to statistically compare distances observed between different segments.

### Confidence intervals and statistics

Error bars, unless otherwise stated, correspond to 95% confidence intervals on the mean. To take a nonparametric approach to determining the statistic, we randomly resampled with replacement from the underlying data distribution 1000 times and calculated the statistic’s bootstrap distribution. Statistical tests are stated throughout the figure legends and text with sample sizes and p values.

## Supplemental Information

Figures S1-S5

## Supplemental Videos

**Supplemental Video 1. Example 2D position decoding of hippocampal activity corresponding to taken and untaken paths ahead.** Related to Figure 2. Note when approaching first choice point, the hippocampal representation extends ahead along the taken path ahead, while when approaching the final choice point, the hippocampal representation extends along the untaken path ahead. Rat position and head direction shown in magenta, most likely decoded position shown in green, decoded position from hippocampal activity shown in heatmap. Video is slowed 8x.

**Supplemental Video 2. Example 2D position decoding of hippocampal activity corresponding to an untaken path behind.** Related to Figure 3. Note that the hippocampal representation extends into the untaken path behind the animal. Rat position and head direction shown in magenta, most likely decoded position shown in green, decoded position from hippocampal activity shown in heatmap. Video is slowed 8x.

**Supplemental Video 3. Example 2D position decoding of hippocampal activity corresponding to a remote patch.** Related to Figure 4. Note that the hippocampal representation corresponds to positions in an alternative patch. Rat position and head direction shown in magenta, most likely decoded position shown in green, decoded position from hippocampal activity shown in heatmap. Video is slowed 8x.

---

**Figure S1.**
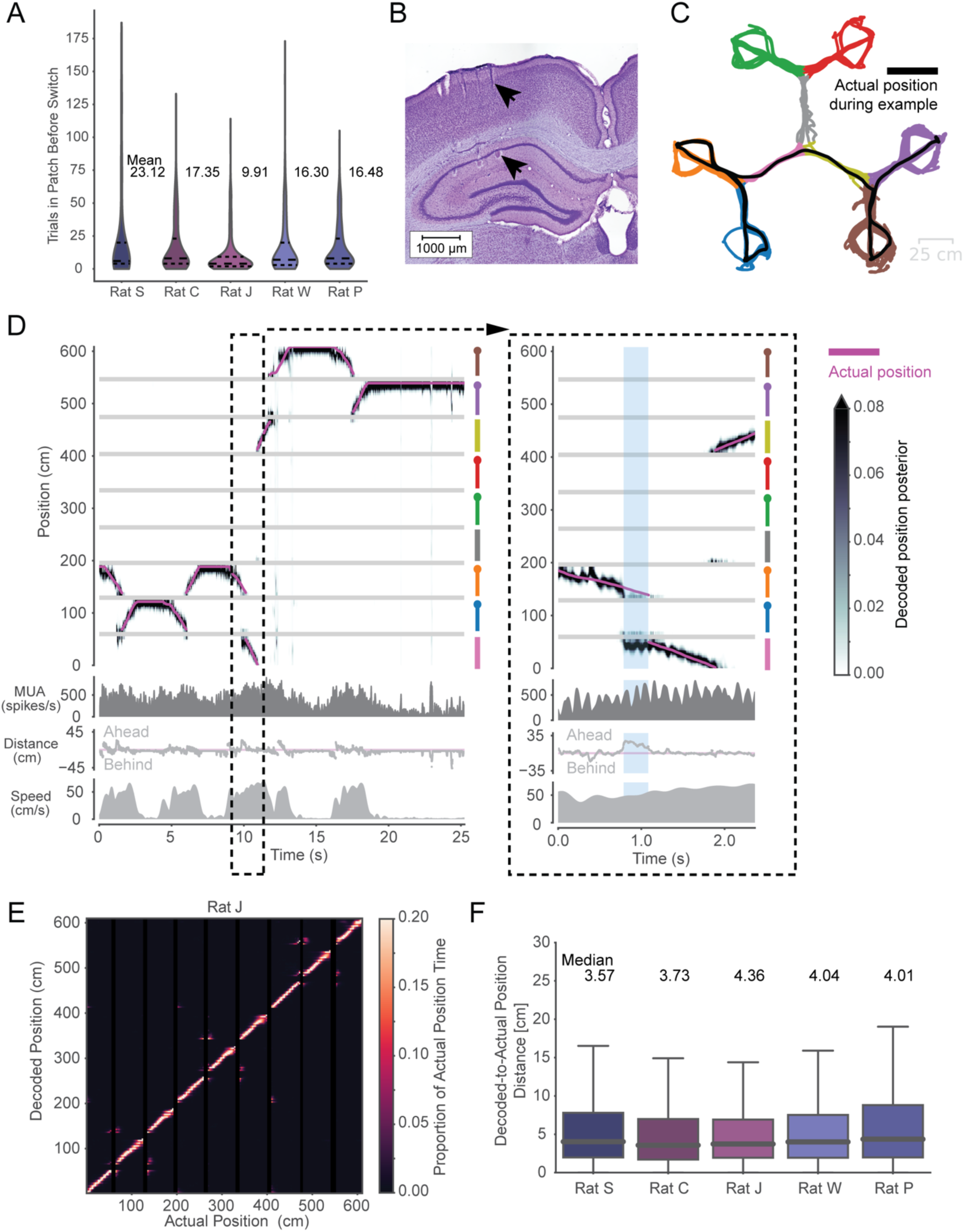
Decoding animal position from Hippocampal spiking during patch foraging. **(A)** Violin plots showing distribution of number of trials spent in a patch before Switching, with horizontal dashed line at median, and upper and lower dashed lines indicating quartiles. Mean trial bout durations are labeled per animal. **(B)** Nissl-labeled coronal brain tissue section representative example showing targeting of tetrodes to dCA1 of the hippocampus. Black arrows point to a tetrode track in dorsal cortex above and a lesion from the tetrode tip in the pyramidal cell layer below. **(C)** Animal head position over the course of one example session, with position data colored by the currently-occupied track segment. Black line highlights the animal’s trajectory throughout the same ∼25s period shown in **D**. Colored track segments correspond to the colored track segments in **D**. **(D)** Decoded position from hippocampal population spiking tracks the animal’s actual position at a behavioral timescale and can sweep ahead or behind the animal to represent non-local positions at a sub-second timescale. Dashed lines highlight a small period that is enlarged at right. This period shows an example non-local representation (blue vertical bars) where the decoded position is in a segment distinct from the animal’s actual position. Top: Actual head position in 1D (linearized) is shown in pink and decoded position posterior is shown in greyscale. Track segments are aligned on the right y-axis, and segment colors correspond to segments in **C**. Circles at the end of colored segment lines represent reward port locations. Note that actual and decoded position are stationary at reward port positions, as animal pokes into reward port. Horizontal grey lines correspond to 15 cm gaps introduced between track segments in linear position space. Top-middle: Multiunit spike rate across hippocampal tetrodes. Note fluctuations at roughly 8 Hz theta frequency, as expected during running. Bottom-middle: Distance of decoded position from animal’s actual position, which can be either ahead (positive values), at (zero cm), or behind (negative values) the actual position. Bottom: Animal head speed. **(E)** Confusion matrix showing actual position and decoded position for one rat. Decoded position largely tracks animal’s actual position across all track segments. Small amounts of off-diagonal density tend to occur at intersections of track segments, corresponding to adjacent positions in 2D space, as expected. **(F)** Boxplots of distribution of distance between decoded and actual position across valid decoded run times for each animal. Boxes show quartiles, whiskers correspond to the data range, and horizontal lines indicate medians, which are also labeled above.

**Figure S2.**
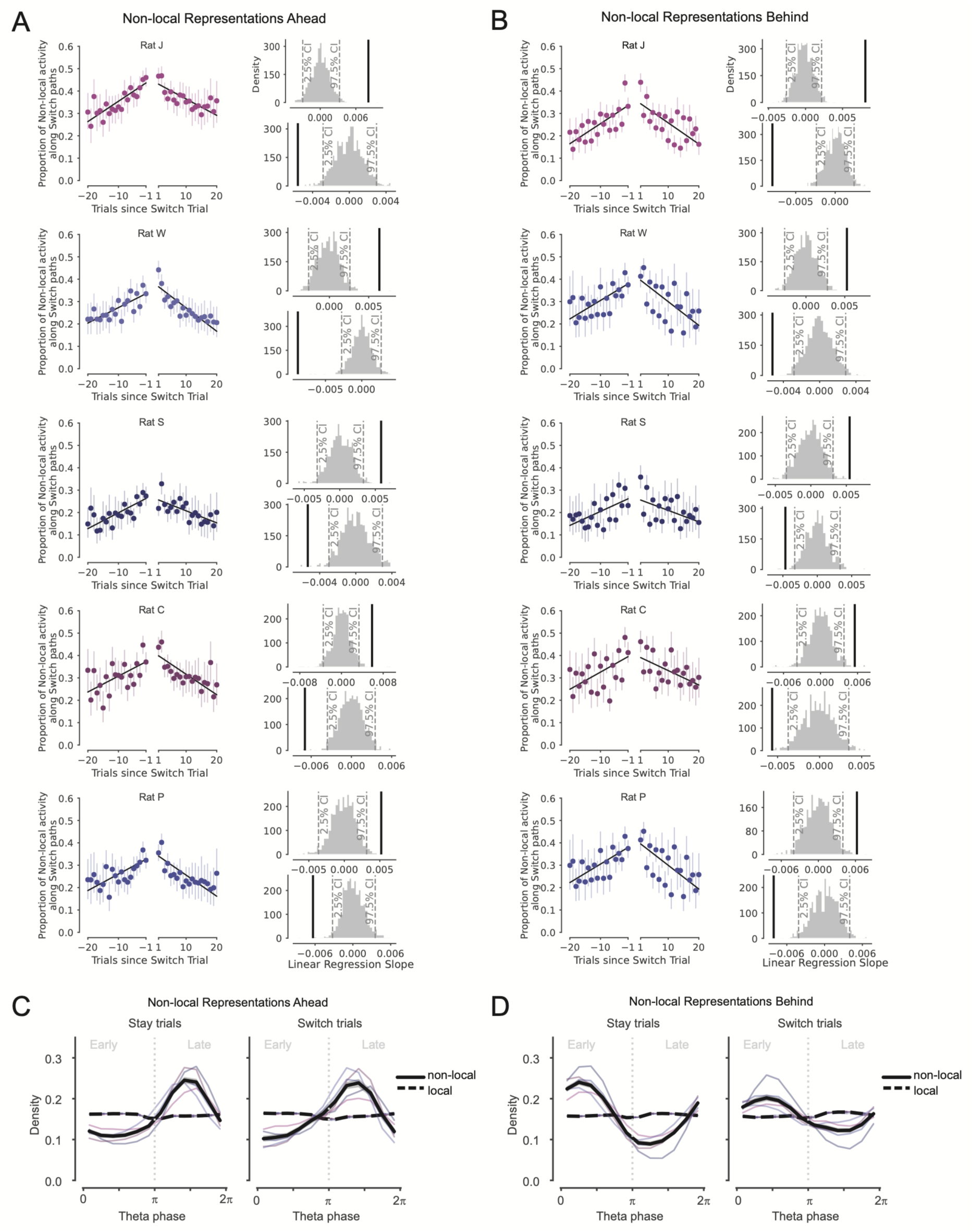
Non-local representations of alternative paths Ahead and Behind are enriched across trials before and after patch Switching. **(A)** Proportion of all non-local activity that represents paths consistent with Switching on Stay trials before and after Switch trial for each animal. Data correspond to Stay trials when animals were located in the first track segment and approaching the choice point, as in Fig. 2B-D. Error bars are 95% CIs on the mean. Pre- and post-Switch linear regressions overlaid in black. All slopes are significantly different than 0 (p_pre_=0.002, 0.002, 0.014, 0.002, 0.004, p_post_=0.002, 0.002, 0.008, 0.006, 0.004). Upper right plot: slope of pre-switch linear regression (black) is greater than the 97.5% CI on the slopes from 1000 shuffles of the underlying data (grey). Lower right plot: slope of post-switch linear regression (black) is less than the 2.5% CI on the slopes from 1000 shuffles of the underlying data (grey). **(B)** Same as **A**, but for non-local activity that represents paths consistent with Switching as the animal traverses the final segment of each trial and approaches the reward port, as in Fig. 3B-D. All slopes are significantly different than 0 (p_pre_=0.002, 0.002, 0.004, 0.01, 0.008, p_post_=0.002, 0.032, 0.004, 0.002, 0.002). **(C)** Non-local representations of paths ahead (solid lines) are concentrated in late phases of the theta rhythm, compared to local representations corresponding to the current track segment (dashed lines), on both Stay trials (left) and Switch trials (right). All animal data in black with 95% CI on the mean in grey band, and individual animal data colored per **A.** Early and late phases are separated with a vertical dotted grey line, and labeled in grey text above. Local and non-local distributions are significantly different in each animal for Stay trials (p = 4.4e-64, 1.0e-16, 2.5e-175,8.2e-129, 2.9e-18, Kuiper test) and Switch trials (p = 1.2e-7, 1.4e-11, 3.3e-12, 3.8-19, 9.3e-14). **(D)** Same as **C**, but for non-local representations of paths behind (solid lines) and local representations corresponding to the current track segment (dashed lines). Non-local paths behind are concentrated in early phases of theta. Local and non-local distributions are significantly different in each animal for Stay trials (p = 2.2e-7, 1.3e-52, 2.9e-202, 3.5e-23, 1.0e-14, Kuiper test) and Switch trials (p = 8.7e-4, 0.040, 1.3e-16, 0.015, 0.016).

**Figure S3.**
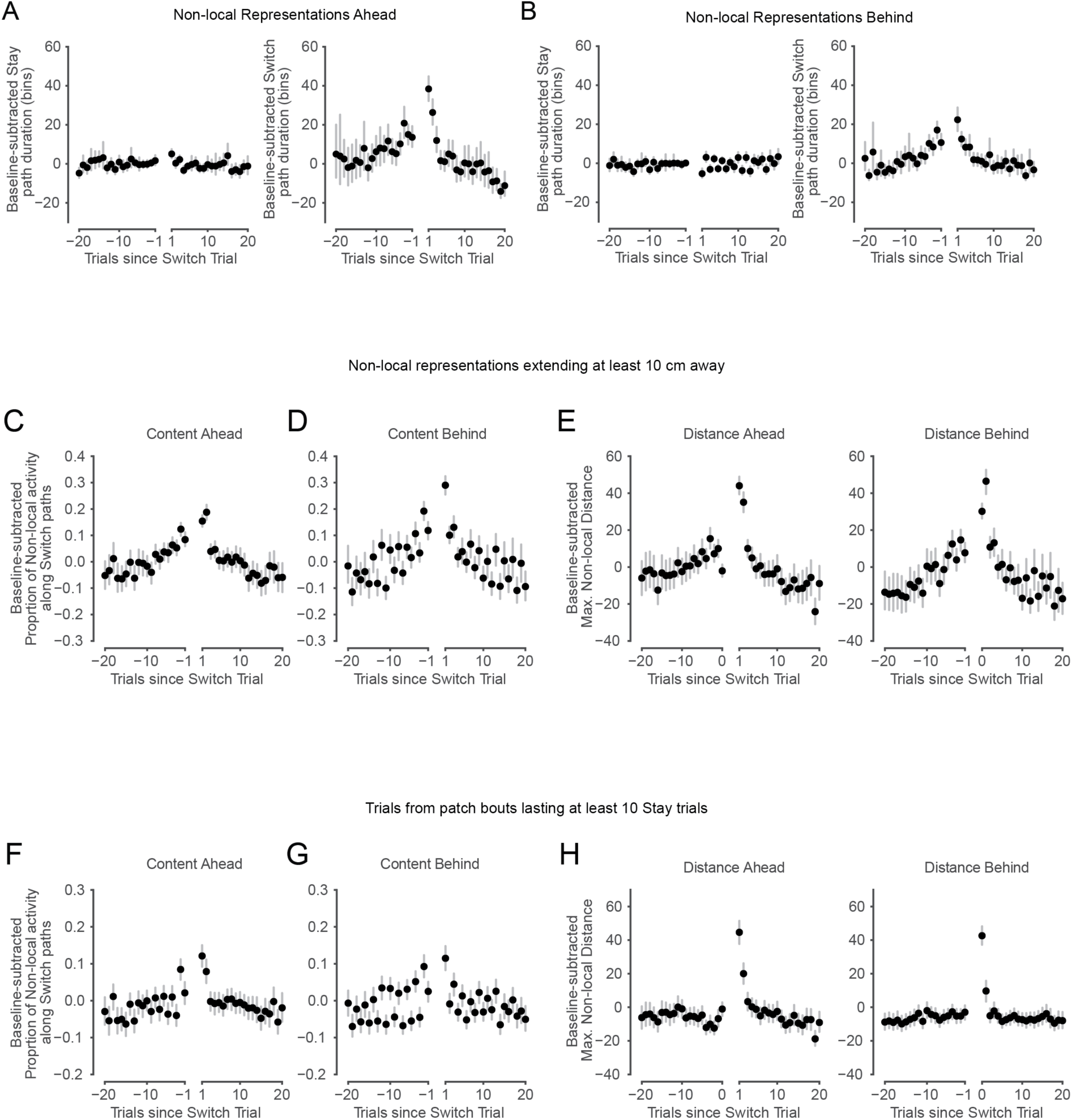
Non-local representations Ahead and Behind are flexibly engaged around Switch trials. **(A)** Baseline-subtracted Stay path duration (left) and Switch path duration (right) across all animals. Durations are in 2 ms bins and quantify the number of bins where non-local representations were present during the approach of the first choice point across trials before and after Switch trials. Error bars are 95% CIs on the mean. Switch path durations were modulated more than were Stay path durations leading up to and following Switch trials. Related to Fig. 2D. Baseline durations per animal ranged 21.12-25.68 bins for Stay path representations and 14.61-28.03 bins for Switch path representations. Non-local representations were detected on 51.1-86.9% of trials per animal. **(B)** Same as **A**, but for non-local representations occurring during traversal of the final track segment and approach of the reward port across trials before and after Switch trials. Related to Fig. 3D. Baseline durations per animal ranged 13.99 – 26.65 bins for Stay path representations and 5.7-14.17 bins for Switch path representations. **(C)** Baseline-subtracted proportion of non-local representations along Switch paths, across all animals. These representations were expressed during the approach of the first choice point across trials before and after Switch trials. Only non-local representations at least 10 cm from the animal are included. Error bars are 95% CIs on the mean. Proportion increases before Switch trials and decreases following Switch trials. Pre- and post-switch linear regression slopes are significantly different than 0 compared to shuffles, as in Fig. S2A (slope_pre_=0.0073, p=0.002; slope_post_=-0.0094, p=0.002). The similarity to Fig. 2D demonstrates that our results are not driven by representations very close to the animal. **(D)** Same as **C**, but for non-local representations occurring during traversal of the final track segment and approach of the reward port. Proportion increases before Switch trials and decreases following Switch trials. Pre- and post-switch linear regression slopes are significantly different than 0 compared to shuffles, as in Fig. S2B (slope_pre_=0.0099, p=0.002; slope_post_=-0.011, p=0.002). The similarity to Fig. 3D demonstrates that our results are not driven by representations very close to the animal. **(E)** Baseline-subtracted maximum non-local distance occurring during approach of the first choice point (left) and traversal of the final track segment and approach of the reward port (right), across all animals. Only non-local representations at least 10 cm from the animal are included. Error bars are 95% CIs on the mean. Distances are especially elevated upon Switching patches and for a couple trials thereafter, and then decrease toward baseline levels. The similarity to Figs. 4D and G demonstrates that our results are not driven by representations very close to the animal. **(F)** Baseline-subtracted proportion of non-local representations along Switch paths, across all animals. Data correspond to the approach of the first choice point across trials before and after Switch trials. Here, non-local representations are only included from trials in bouts within a patch lasting at least 10 Stay trials. Error bars are 95% CIs on the mean. Proportion increases before Switch trials and decreases following Switch trials. Pre- and post-switch linear regression slopes are significantly different than 0 compared to shuffles, as in Fig. S2A (slope_pre_=0.0033, p=0.002; slope_post_=-0.0047, p=0.002). The approximately symmetrical pattern around the Switch is similar to that seen in Fig. 2D, indicating that the results are not only driven by periods where animals Stayed in a patch for a small number of trials. **(G)** Same as **F**, but for non-local representations occurring during traversal of the final track segment and approach of the reward port across trials before and after Switch trials. Error bars are 95% CIs on the mean. Pre- and post-switch linear regression slopes are significantly different than 0 compared to shuffles, as in Fig. S2B (slope_pre_=0.0034, p=0.002; slope_post_=-0.0036, p=0.002). The approximately symmetrical pattern around the Switch is similar to that seen in Fig. 3D, indicating that the results are not only driven by periods where animals Stayed in a patch for a small number of trials. **(H)** Baseline-subtracted maximum non-local distance during the approach of the first choice point (left) and traversal of the final track segment and approach of the reward port (right) across all animals. Here, non-local representations are only included from trials in bouts within a patch lasting at least 10 Stay trials. Distances are asymmetric around Switch trials, and are especially enhanced upon Switching patches, and then decrease across trials after the Switch. Related to Figs. 4D and G, respectively.

**Figure S4.**
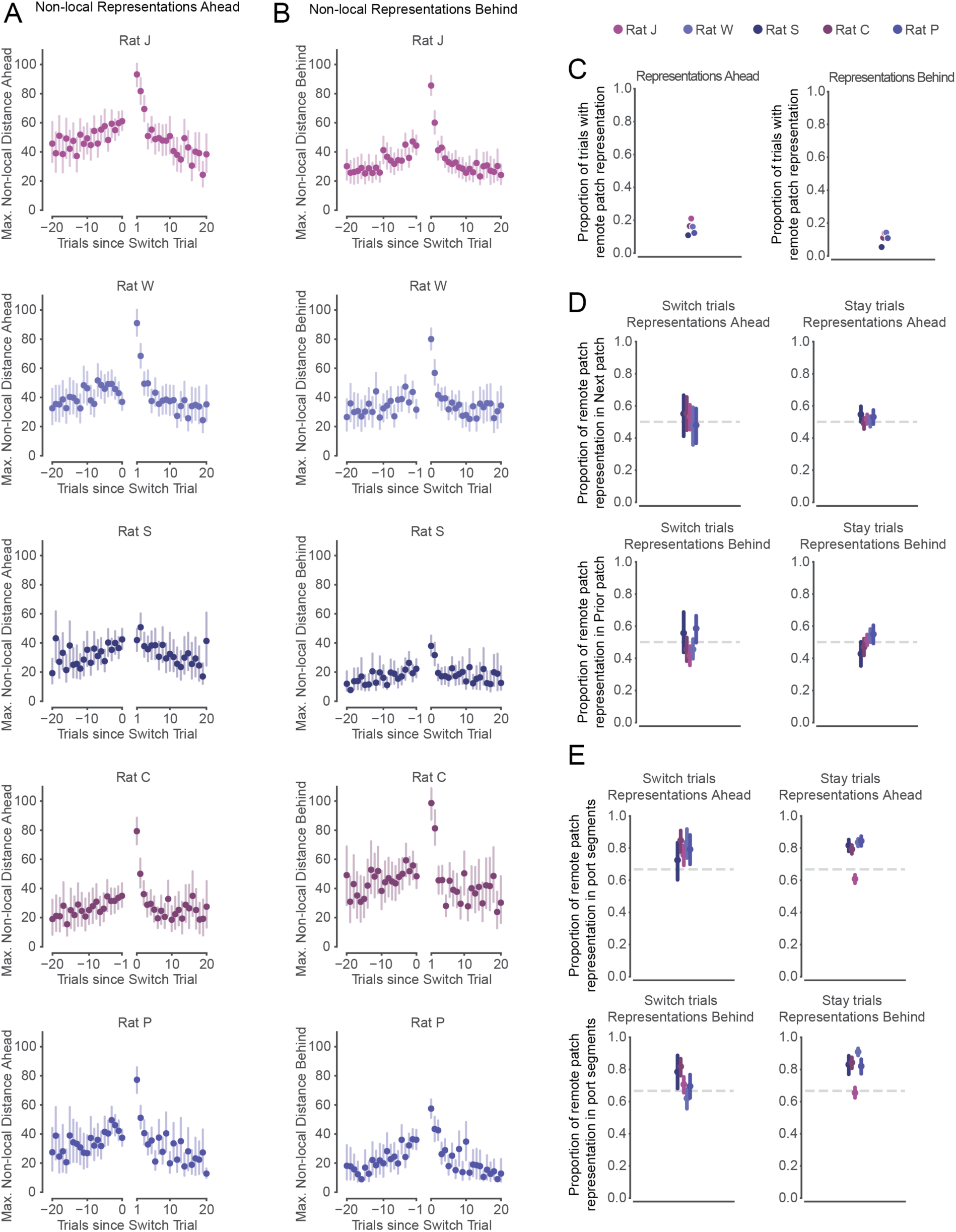
Non-local representations of distant locations. **(A)** Maximum non-local distance represented on trials leading up to and following patch Switches for each animal. Data are from the period of each trial in which the animal approached the first choice point. Error bars are 95% CIs on the mean. Related to Fig. 4D. **(B)** Maximum non-local distance represented on trials leading up to and following patch Switches, during the period of each trial in which the animal traversed the final track segment towards the reward port, for each animal. Error bars are 95% CIs on the mean. Related to Fig. 4G. **(C)** Proportion of trials with non-local representations corresponding to remote, unoccupied patches, out of all trials with any non-local representations, for each animal. Left: calculated for non-local representations occurring as animals approached first choice point. Right: calculated for non-local representations occurring as animals traversed final track segment and approached reward port. In both cases, ∼10-20% of trials contain representations corresponding to locations in remote patches. Note that trials with any non-local representations corresponded to 76% of all trials. **(D)** Non-local representations corresponding to locations in remote patches are not biased to represent subsequently chosen or immediately previous patches. Top row: proportion of remote patch representations occurring during approach of first choice point on Switch trials (left) and Stay trials (right) that correspond to subsequently chosen (next) patch. Bottom row: proportion of remote patch representations occurring during traversal of final track segment and approach of reward port on Switch trials (left) and Stay trials (right) that correspond to most recently chosen (prior) patch. Error bars are 95% CIs on the mean. Grey dashed lines correspond to an equal (50%) representation of the chosen and non-chosen patches. **(E)** Non-local representations in remote patches are approximately equally distributed across track segments. Top row: proportion of remote patch representations occurring during approach of first choice point on Switch trials (left) and Stay trials (right) that correspond to track segments containing reward ports. Bottom row: proportion of remote patch representations occurring during traversal of final track segment and approach of reward port on Switch trials (left) and Stay trials (right) that correspond to track segments containing reward ports. Error bars are 95% CIs on the mean. Grey dashed lines correspond to a chance (66.67%) representation of the 4 segments containing reward ports out of 6 total track segments in remote patches.

**Figure S5.**
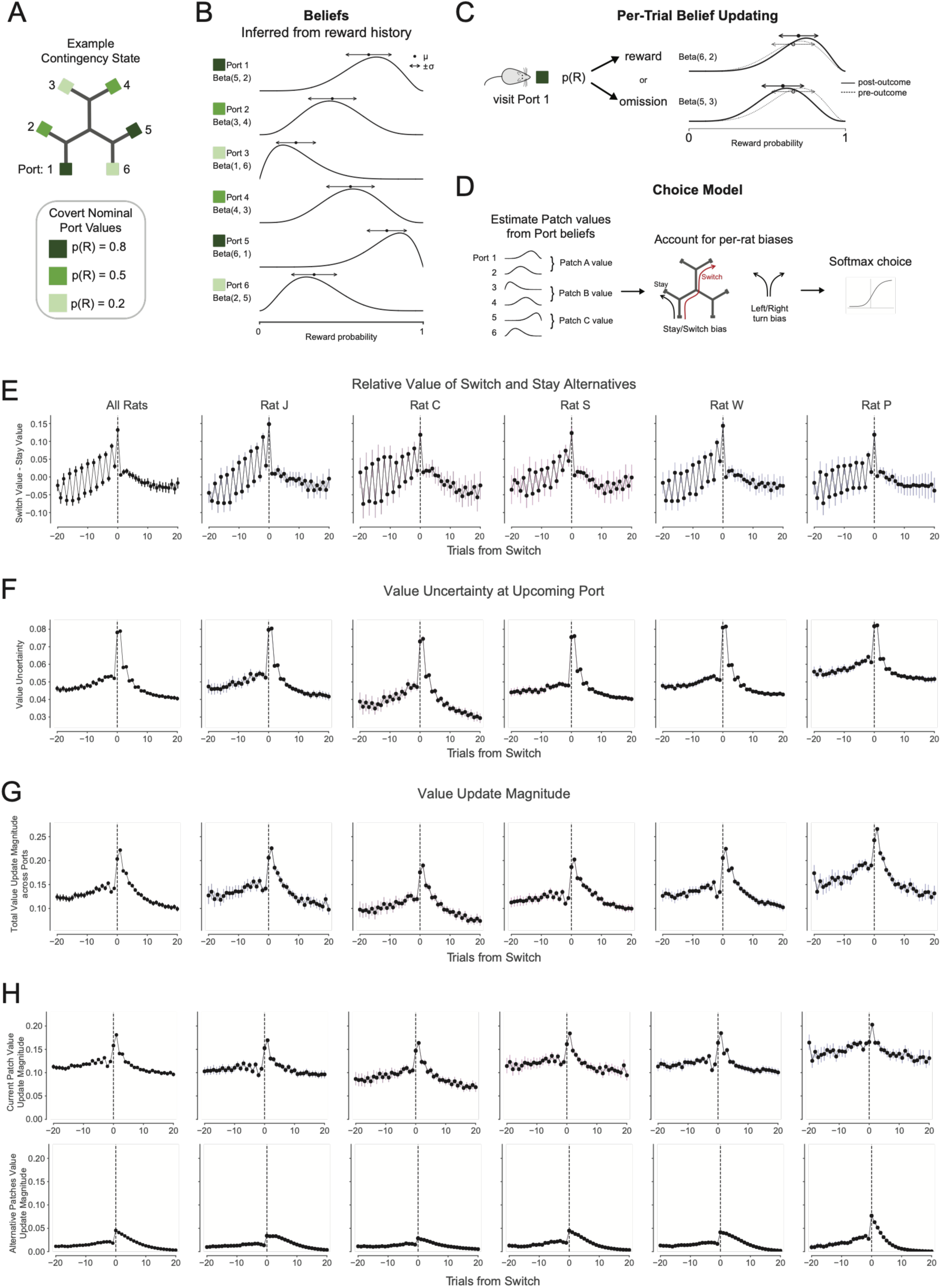
Beta-Bernoulli behavioral model schematic. **(A)** Schematic of an example reward contingency, with each port assigned a hidden nominal reward probability p(R). **(B)** Schematic of beliefs of the value of each Port 1-6. Beliefs are based on counts of binary reward and omission outcomes, which are observed over trials from each independent Bernoulli port. The model maintains a Beta posterior distribution over the reward probability of each port. These schematized distributions correspond to seven simulated visits per port, with the number of rewards and omissions in the labeled Beta function; for example, Port 1 with Beta(5,2) illustrates beliefs after five rewards and two omissions. Port-level beliefs on each trial can be represented by a mean and variance, μ and σ^2^, which correspond to an expected value and uncertainty. **(C)** On each trial, either a reward or omission is observed, which updates the visited port’s value belief distribution accordingly. The schematized example shows a visit to Port 1, where the pre-outcome beliefs (dotted line) are as in B after seven simulated visits, and the post-outcome beliefs (solid line) show the updated distribution, which shifted higher after reward (upper plot), or shifted lower after omission (lower plot) on a simulated eighth visit. Note that learning is local; each outcome yields an updated mean and variance of the visited port’s beliefs. The un-visited port beliefs are not updated based on the outcome; only a small relaxation towards a uniform distribution is allowed to account for the possibility of forgetting. **(D)** To connect the value learning model in **C** to choices and enable us to assess the model’s ability to capture animal behavior, we computed choice likelihoods on each trial. Left: We estimate Patch values from averaged beliefs about the mean values of the two ports within each Patch (left). See methods for details. Middle: Bias terms, fit per-animal, are then integrated into the inferred Patch values. Two example biases are illustrated: a Stay/Switch bias, sensitive to the cost of Switching to a more distant port, and a Left/Right turn bias. Right: The likelihoods of patch choices on each trial are computed as the softmax of the three patch values. On Switch trials, likelihood of port choice is computed with a subsequent softmax between the values of the two ports in the new patch. **(E)** Relative value of Switching and Staying for each animal, estimated with the behavioral model based on animal’s experienced reward histories. The relative value of Switching increases on average leading up to Switch trials and decreases afterwards. Error bars are 95% CIs on the mean. Related to Fig. 1F. **(F)** Uncertainty around the model-estimated value of the upcoming (chosen) port on each trial, for each animal, reflecting variances of value beliefs schematized in Fig. S5B above. Uncertainty is elevated on and after patch Switches. Across-trial dynamics are asymmetric around the Switch. Error bars are 95% CIs on the mean. Related to Fig. 5B. **(G)** Value updates, measured as the absolute magnitude of the change in model-estimated values across ports, for each animal. The degree of value updating is elevated on and after patch Switch trials, then decays across trials back to baseline. Error bars are 95% CIs on the mean. Related to Fig. 5B. **(H)** Value updates as in **G**, separated into updates of beliefs about the currently chosen Patch (top) and updates about the alternative unchosen patches (bottom). Note distinct magnitudes; while chosen port values are updated by reward outcomes, small changes to values of unvisited ports occur only as a result of “forgetting” over time since last visit. Error bars are 95% CIs on the mean. Related to Fig. 5B.

